# TreeKnit: Inferring Ancestral Reassortment Graphs of influenza viruses

**DOI:** 10.1101/2021.12.20.473456

**Authors:** Pierre Barrat-Charlaix, Timothy G. Vaughan, Richard A. Neher

**Affiliations:** Biozentrum, Universität Basel, Switzerland; Swiss Institute of Bioinformatics, Switzerland; ETH Zurich, Department of Biosystems Science and Engineering, Basel, Switzerland

## Abstract

When two influenza viruses co-infect the same cell, they can exchange genome segments in a process known as reassortment. Reassortment is an important source of genetic diversity and is known to have been involved in the emergence of most pandemic influenza strains. However, because of the difficulty in identifying reassortments events from viral sequence data, little is known about its role in the evolution of the seasonal influenza viruses. Here we introduce TreeKnit, a method that infers ancestral reassortment graphs (ARG) from two segment trees. It is based on topological differences between trees, and proceeds in a greedy fashion by finding regions that are compatible in the two trees. Using simulated genealogies with reassortments, we show that TreeKnit performs well in a wide range of settings and that it is as accurate as a more principled bayesian method, while being orders of magnitude faster. Finally, we show that it is possible to use the inferred ARG to better resolve segment trees and to construct more informative visualizations of reassortments.

## I. Introduction

Influenza viruses evolve rapidly and change their surface proteins, which allows them to evade preexisting immunity and reinfect their hosts. The viral genome is made of 8 RNA segments that encode for 11 different proteins, with segments coding for the surface proteins haemagglutinin (HA) and neuraminidase (NA) being the most important for immune escape. In each segment, evolution is an asexual process in which diversity is generated by mutations. However, when a host cell is simultaneously infected by more than one virus, offspring viruses can carry segments from several parents – a process know as reassortment. Genomic reassortment is akin to sexual reproduction and can generate viruses with novel genetic constellations. In particular, it has been found to be the cause of most pandemic influenza strains [1, 2].

The genealogy of a single segment is described by a tree, whose leaves correspond to observed sequences and internal nodes to the ancestry of different lineages. Many methods exist to reconstruct this tree from gene sequences [3–5]. However, trees are not well suited to describe genealogies of full genomes in the presence of reassortment since lineages can then have difference ancestors for their different segments. A more adapted concept is the so-called Ancestral Reassortment Graph (ARG), or Ancestral Recombination Graph in the context of recombination. Internal nodes of the ARG represent either coalescence of different lineages, in which case they have a unique ancestor as internal tree nodes, or the emergence of a new lineage from a reassortment, in which case they may have several ancestors. Simple examples of ARGs for two segments are shown in Figure 1.

**Figure 1.**
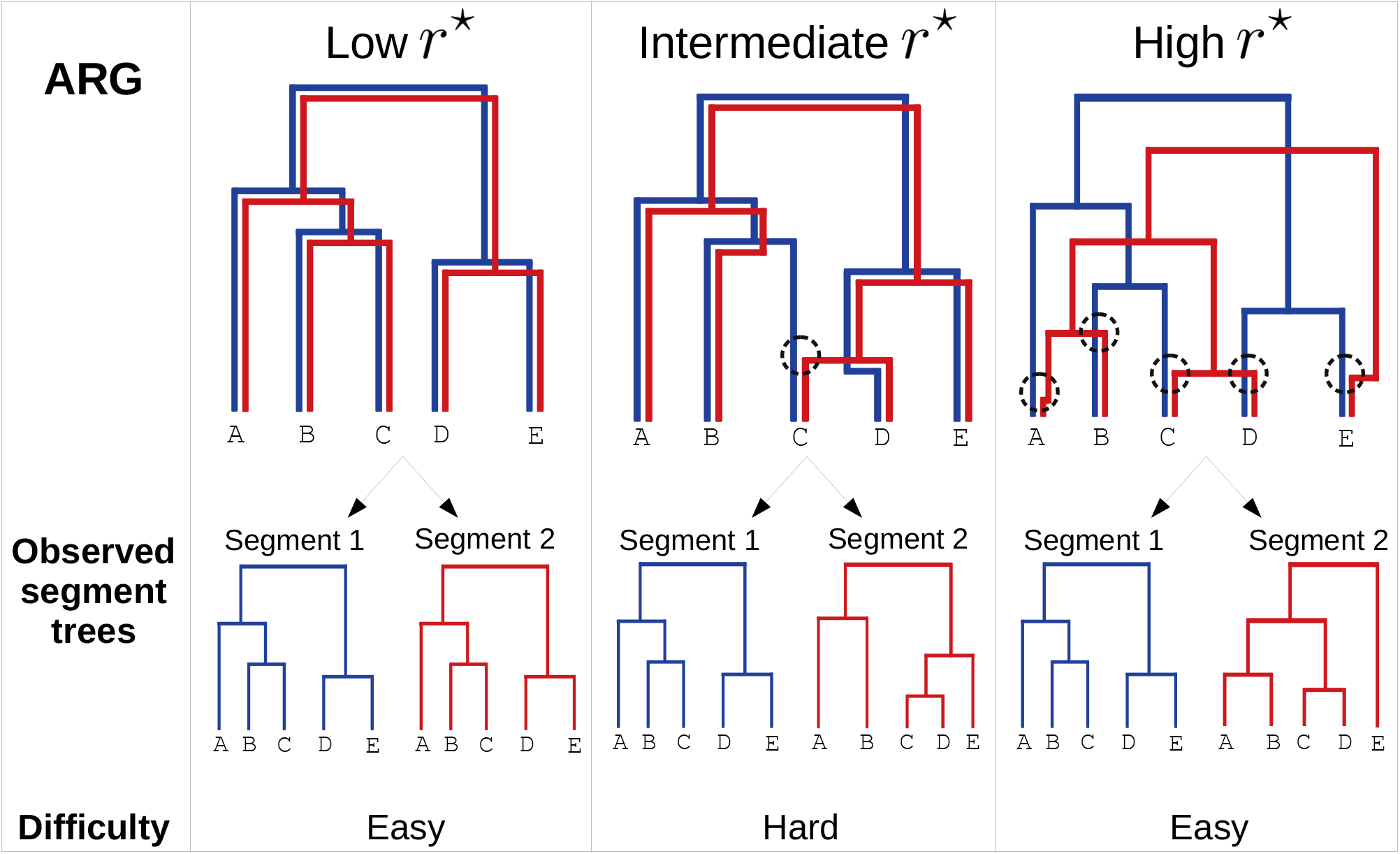
Example of ARGs for five sampled strains and for two segments (blue and red). Reassortments are shown as black circles in each ARG. Based on the scaled reassortment rate in the population *r*^*\**^, three regimes can be identified. **Left**: Very low reassortment rate. Reassortments are very rare, and every strain inherits its two segments from the same parent. The ARG is equal to the gene trees, and reconstructing it is easy. **Center**: Intermediate reassortment rate. Some exchange of segments takes place: some strains do not inherit their segments from the same parent (here, strain `C`). The segments trees have different topologies, but are still relatively similar. Inferring the position of reassortments from the gene trees is non trivial. **Right**: Very high reassortment rate. A reassortment takes place on every branch of the ARG before the first coalescence. The two segments have independent evolutionary histories, and the segment trees share no structure. Inference of reassortments becomes easy again.

The knowledge of the ARG for a set of influenza sequences would be of major interest, mainly as it would shine light on the role of reassortment in the evolution of influenza. While the importance of major reassortment events in the formation of pandemic strains is known, much less can be said on the effects of smaller scale intra flu-lineage reassortments. Several studies have tackled this problem, with significant discrepancy in their results [6–8]. This is likely due to the lack of a robust and efficient method to infer the whole set of reassortments in the history of a large and representative set of viral genomes. In addition, knowing the ARG would allow a more accurate reconstruction of tree branch lengths in regions without reassortments by using the sequences of several segments, or a better visualization of pairs of trees of different segments by disentangling tanglegrams as much as possible (see Figure S9 in SI).

A common method to identify reassortments is to reconstruct segment trees and manually compare them [9, 10], which is time consuming and error prone. A number of automated methods have also been developed. Some do not go through the step of reconstructing phylogenies and instead compare the sequence distance of different strains for different segments [11, 12]. Other approaches consist in finding discrepancies between segment trees, either using topology [13–15] or mutation patterns on the branches of the trees [7]. Sets of probable reassortments are then deduced from these discrepancies, typically using a confidence score. A common point to all these methods is that they only identify a subset of the reassortments that occurred in the genealogy, which could result in a mis-interpretation of the importance of reassortment. Additionally, it is not possible to fully reconstruct the genealogy since differences between trees remain even after accounting for the inferred reassortments. Recently, a method for directly inferring the ARG from sequences has been proposed that extends the principles used to infer phylogenetic trees to data containing reassortments [8, 16]. It does so by using a coalescence-reassortment model to assign a probability to any ARG based on observed sequences and then samples from this probability. While this model based approach is appealing, it is computationally expensive and limited to medium size datasets.

Here we propose TreeKnit, a fast method to infer ARGs from pairs of segment trees that processes by “knitting” the trees together starting from the leaves. The underlying idea is that topological differences between trees are caused by reassortments, and that we can thus introduce reassortments so as to minimize these incompatibilities. In a first part, we describe how TreeKnit works. We then estimate its performance and limitations on simulated genealogies and show that it can be used to better resolve segment trees. Finally, we compare it to two existing methods to infer reassortments in influenza genealogies.

## II. Methods

Whether inferring ARGs is easy or hard and whether it is useful or not depends on the relative strength of coalescence and reassortment. Qualitatively, we can distinguish three main regimes represented in Figure 1. For a very low reassortment rate, reassortments are so rare that the ARG can be considered tree-like, with the two segment trees being identical. Recovering the ARG from the knowledge of the segment trees is then trivial. On the contrary, for a very large reassortment rate, the first reassortments along a lineage occur well before any pair of strains coalesce to a common ancestor. The two segments evolve in practice independently, and their trees have no shared structure. Inferring the ARG is again easy although uninformative: one only has to introduce a reassortment above each leaf. The intermediate case is both the hardest and the most interesting one. Indeed, reassortments are then rare enough that the segment trees share a lot of structure, but sufficiently frequent for the problem to not be trivial.

### Maximally Compatible Clades (MCC)

A central concept for our method are *Maximally Compatible Clades (MCCs)*. Given a two segment ARG that embeds two trees, one obtains the MCCs by removing all branches that correspond to only one tree, and keeping those that are common to both. In Figure 1, this would amount to removing all branches that have only one color and keeping those that are both red and blue. This operation results in a set of disjoint trees, each of those being one MCC. Note that MCCs are not necessarily clades in the segment trees, since they can be nested. It is convenient to refer to an MCC by the leaves that it contains, and we will do so in the following.

MCCs have several properties that makes them a very useful concept for thinking about ARGs: (i) If both segment trees *and* all MCCs are known, so is the observable part of the corresponding ARG. This follows from the fact that MCCs are the regions where the two trees are “stitched” together in the ancestral graph. Our method reconstructs ARGs using this idea and is effectively a method to *infer MCCs given a pair of trees*. Once the MCCs are known, the only information missing to fully reconstruct the ARG are the times at which reassortments occurred on internal branches. (ii) There is a one-to-one correspondence between MCCs and observable reassortment events. This is a simple consequence of the fact that reassortments correspond to the separation of lineages of different segments in the ARG, and therefore to the transition between a region where branches are common to both trees, and a region where they are not. By definition the root of an MCC must be located right where this separation occurs. The only exception to this rule is the case of an MCC that contains the root one of the segment trees. The implication is that the number of reassortments in the genealogy of two segments is equal to the number of MCCs in their ARG, minus one if one of the MCCs contains a root. (iii) Restricting segment trees to an MCC results in two subtrees with the same topology. MCCs are maximal in the sense that extending them by adding nodes results in *topologically different* subtrees in the two segments. This last property is important for our method: finding MCCs is equivalent to finding maximal sets of leaves that give rise to subtrees with matching topologies.

### Finding MCCs: the TreeKnit method

Given a set of compatible clades (CCs), initially the leaves, TreeKnit will attempt to grow them until they are maximal by cycling through four steps illustrated in Figure 2 and detailed below:

**Figure 2.**
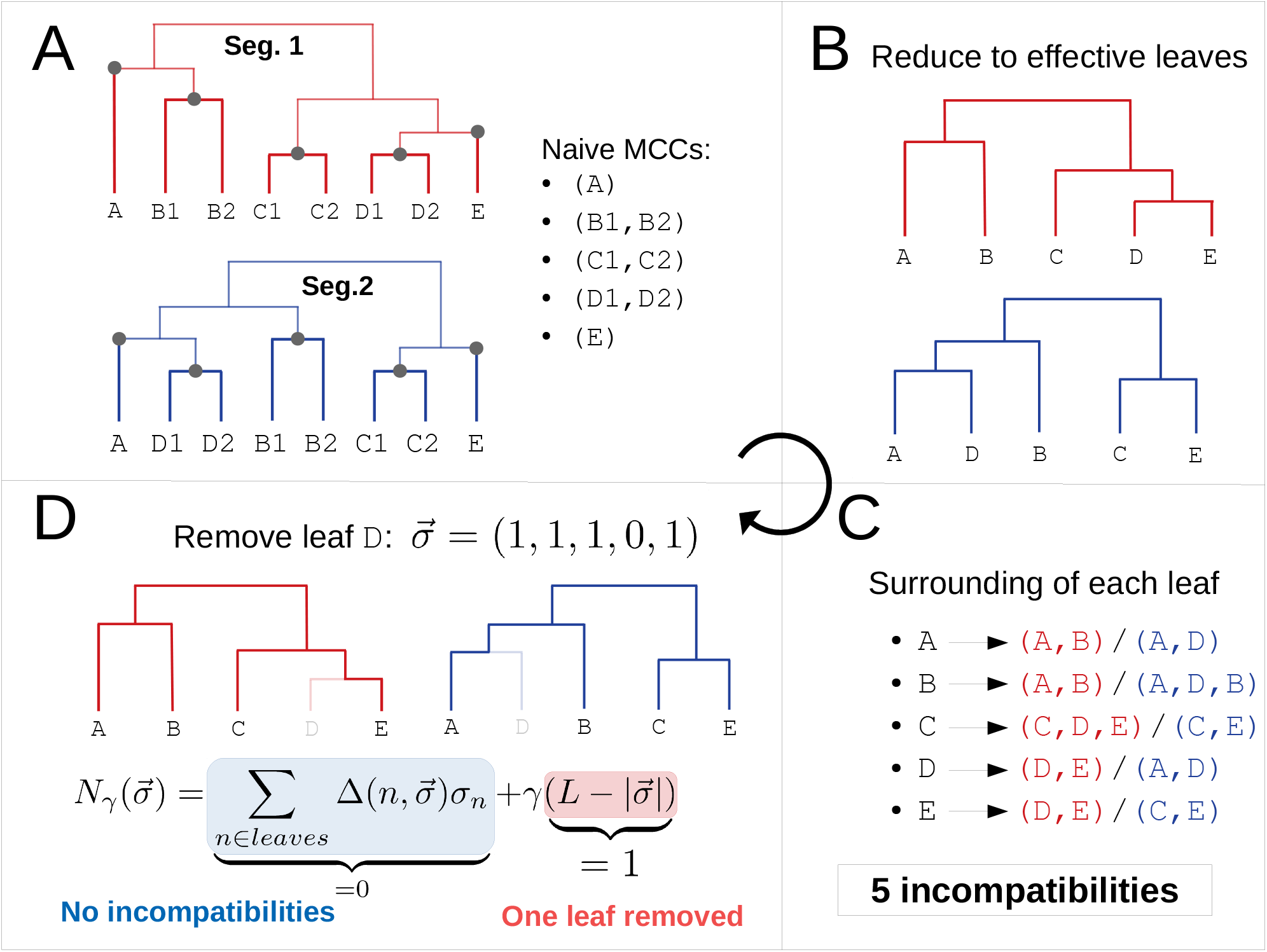
Schematic of the iterative algorithm. **A**: Construction of the naive MCCs. Circles indicate the root of the five clades that match exactly in the two trees (slightly highlighted branches). Trying to grow one of these clades gives inconsistent results in the two trees: *e*.*g*. growing the MCC `(B1,B2)` gives clade `(A,B1,B2)` in the first tree and `(A,D1,D2,B1,B2)` in the second. **B**: Trees obtained after reducing trees of **A** to their naive MCCs: each clade is represented by a single effective leaf. **C**: Counting incompatibilities in the reduced trees. For each effective leaf, the clades defined by its direct ancestor in the two trees are compared, and each mismatch counts as one incompatibility. **D**: Enforcing reassortments on some leaves to remove incompatibilities. A configuration 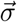 is associated to each set of removed leaves. The scoring function 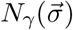 adds the number of remaining incompatibilities given 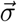 and the number of removed leaves multiplied by *γ*. The optimal set of reassortments is found by minimizing 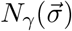, *e*.*g*. removing `D` is optimal if *γ <* 5.

i. Perform a naive maximization of compatible clades (CCs): grow CCs by adding internal nodes as long as the obtained clades are exactly equivalent in the two trees (Figure 2**A**). The resulting compatible regions are called the naive MCCs, where the “maximal” term will be justified in the next section.
ii. Collapse the naive MCCs into effective nodes (Figure 2**B**). This allows us to ignore the topological details of the putative CCs. Note that if we applied step *(i)* to the reduced trees, we would find naive MCCs consisting only of leaves, by construction.
iii. Count topological incompatibilities in the reduced trees (Figure 2**C**). For each leaf, compare its surroundings in the two trees, and count one incompatibility if the two surroundings do not match. The surrounding of a leaf is defined as the clade defined by its parent node.
iv. Enforce reassortments on some leaves in order to *minimize* the number of incompatibilities (Figure 2**D**). Effective leaves above which a reassortment is enforced are *removed from the trees*, which reduces the number of incompatibilities. The cost associated with removing a leaf is determined by parameter *γ*. Effective leaves removed at the end of this operation are added to the list of final MCCs. If no leaf was removed as a result of the optimization, *i*.*e*. if no reassortment was enforced, stop here. Otherwise, go back to step *(i)*

A leaf *n* can give rise to an incompatibility counted in step *(ii)* for two reasons: If reassortment took place on the branch leading to *n*, it will be observed in different regions in the two trees, corresponding to the two ancestral viruses that took part in the reassortment. Or *n* is “close” to another reassorted leaf in one of the trees and not in the other, in which case it will also have different surroundings. In the example of Figure 2, `D` corresponds to the first situation, and `A,B,C` and `E` to second.

Given two reduced trees of *L* effective leaves after step *(iii)*, the idea of step *(iv)* is to reduce as much as possible the number of topological incompatibilities by removing (mark as reassortants) a minimal number of leaves. We encode the current state of each leaf *n* in the binary variable *σ*_*n*_ ∈ {0, 1}: it is set to 1 if leaf *n* is present in the tree, and to 0 if it is removed from the tree. Furthermore, define 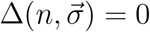 if the trial configuration 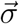 resolves the incompatibility above node *n* (compare Figure 2**C&D**), and 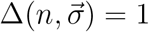 otherwise. For any combination of removed leaves 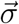, we now define the scoring function 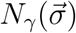:

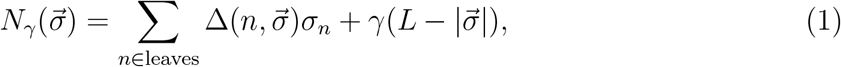

where 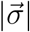 is the *l*1-norm of 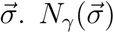 is composed of two terms with a simple interpretation: the first sums over leaves and counts the number of remaining incompatibilities, while the second counts the number of removed leaves with a removal cost *γ*.

The optimal set of leaves to remove is then found by finding the configuration 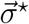 that minimizes *N*_*γ*_. Minimization is performed efficiently by simulated annealing [17], see SI. In the case of Figure 2**D** and for *γ* ≤ 5, the optimal configuration is the one that removes leaf `D` only, that is 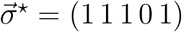, with a score 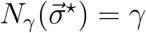. Removing any single other leaf gives a score 4 + *γ*, and is thus always worse than removing `D`. Not removing any leaf gives a score 5, which becomes optimal if *γ >* 5.

The parameter *γ* plays a key role in the result of the method and thus on the reconstruction of the ARG. In the next paragraphs, we discuss the behavior of TreeKnit for extreme values of *γ* and explain some underlying ideas.

### Naive MCCs

*γ* → ∞ If *γ* is very large, *i*.*e*. of the order of the number of leaves in the reduced trees, removing leaves in step *(iv)* has a prohibitive cost. The MCCs returned by the algorithm will thus be the ones found in step *(i)*, that is the *naive MCCs*.

This *n*aive approach has a simple interpretation: any two clades with the exact same topology in two trees are matched, but compatible regions are not extended further. The limit can thus be thought of as a *conservative* approach to the reconstruction of MCCs that avoid over-extending MCCs.

On the other hand, since there is a one to one mapping between MCCs and inferred reassortments, the naive method tends to over-estimate the number of reassortments. This evident in Figure 2: five naive MCCs are found, corresponding to five inferred reassortments since none of them contains the root of one of the trees. The naive method introduces a reassortment for each incompatibility it finds and treats the two types of incompatibilities discussed above identically. However, to infer a more accurate set of MCCs, it is necessary to identify the first type of incompatibility corresponding to reassortment along the branch leading to the leaf. After identifying these and removing the corresponding MCC, other incompatibilities tend to resolve as is the case in the example in Figure 2: A single reassortment above clade `(D1, D2)` explains the two trees.

### Approximately parsimonious MCCs

*γ* = 1 We have seen above that every incompatibility gives rise to one inferred reassortment in the naive approach. On the other hand, we also know that an incompatibility above one leaf might not be due to a reassortment above it, but rather to its proximity to another reassorted leaf. This means that if we observe *N* incompatibilities, we can often resolve them by using less than *N* reassortments.

For *γ* = 1, the scoring function *N*_*γ*_ of Equation 1 has a simple interpretation in terms of *parsimony*. Its first term counts the number of remaining incompatibilities, approximating the number of remaining reassortments predicted by the naive approach once the leaves with *σ*_*n*_ = 0 have been removed. The other term 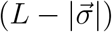 counts the number of removed leaves, which are *enforced* reassortments. 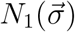 thus approximates the total number of reassortments for a configuration 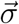. Since the configuration 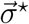 minimizes *N*_1_, it can be interpreted as a parsimonious explanation of the topological differences between the trees.

### Bridging parsimonious and naive approaches

**intermediate** *γ* Figure 2 shows an example where a parsimonious method clearly outperforms the naive one. In some cases, for example the high reassortment rate case shown on the right panel of Figure 1, this is not true: in this case the correct MCCs consist of single leaves, and the naive approach then performs well. However, it is possible to explain the tree with fewer reassortments. Removing leaves `A,B` and `C`, corresponding to configuration 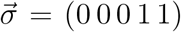, will result in 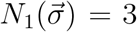, since the two remaining leaves `D` and `E` will then form a compatible clade in the two trees (three enforced reassortments, no incompatibility left). More generally, given an ARG of *L* leaves and with an infinitely high reassortment rate, it is always possible for the pseudo-parsimonious approach to enforce reassortments on *L* − 2 leaves, and have the remaining 2 form a compatible clade, thus obtaining *L* − 2 reassortments in total, instead of *L*. This is not surprising, as it is expected that a parsimonious method performs poorly when there are many reassortments.

Parameter *γ* in Equation 1 tunes the “aggressiveness” with which the algorithm tries to grow compatible clades. In the pseudo-parsimonious approach, *N*_*γ*_ stays constant if one enforced reassortment “fixes” exactly one incompatibility. On the other hand, if *γ >* 1, every removed leaf must fix more than one incompatibility for *N*_*γ*_ to stay constant. As a consequence, it is harder to remove leaves and thus to grow MCCs. In the extreme limit of *γ* → ∞, MCCs cannot be grown and we fall back to the naive method.

The parameter *γ* thus allows us to interpolate between pseudo-parsimonious and naive approaches, and can be thought of as how “conservative” the inference of MCCs is. Note that given the discrete nature of Equation (1), the sharpest changes of behavior of *N*_*γ*_ happen when *γ* crosses an integer value. In the following, we will mostly use integer values *γ*.

### Poorly resolved trees

Trees inferred from sequences are often not completely resolved: branches in the actual genealogy along which no mutations happened will not appear in the reconstructed tree. This results in polytomies or multifurcations at internal nodes with more than two offspring. On a branch of the ARG shared by the two segment trees, it is possible that mutations occurred in one of the segments and not in the other such that a polytomy will be present in one reconstructed segment tree and not in the other. Figure 4**A** gives an example of such a situation.

**Figure 4.**
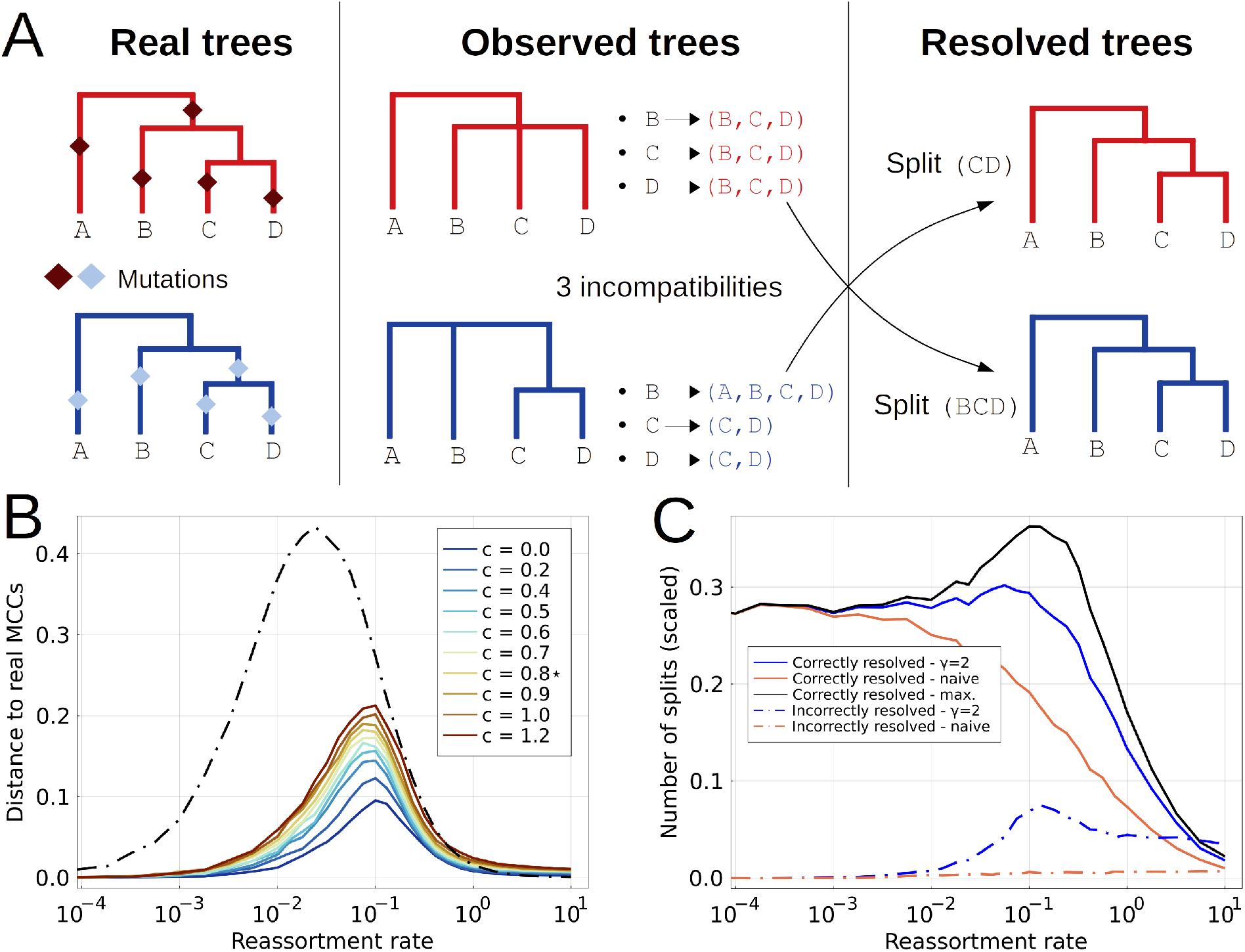
Effect of poorly resolved trees on the inference of MCCs. **A**: Pre-resolving trees before inferring MCCs. The approach is greedy: every split of one tree that is compatible with the other is introduced in the other. **B**: VI distance to real MCCs as a function of *r*^*\**^ for different tree resolutions *c*. The quality of the inference decreases with *c. c*^*\**^ ≃ 0.8 corresponds to levels found in A/H3N2 influenza trees with strains from the same season. **C**: Quality of the resolution of trees *after* having inferred MCCs. This combines splits introduced by the pre-resolution step, and splits known once the MCCs are inferred. The number of correctly inferred splits is shown, scaled by the number of splits that would be necessary to make the trees binary. The black line indicates the performance if the MCCs were exactly known.

If the polytomies are in parts of the segment trees shared in the ARG, one can improve phylogenetic resolution by using the sequences of both segments to infer topology and branch length. However, these shared parts of the ARG need to be identified first, which in turn is hampered by lack of phylogenetic resolution.

In order to overcome this problem, it is necessary to disentangle topological incompatibilities that are due to reassortment from those due to lack of resolution. We reduce the latter by resolving polytomies in each tree using the clades observed in the other. Formally, given two trees 𝒯_1_ and 𝒯_2_, we introduce in 𝒯_1_ every split of 𝒯_2_ that is compatible with the set of splits in 𝒯_1_, and inversely. This assumes that topological differences that could be trivially explained by lack of phylogenetic resolution are never due to reassortment.

The right panel of Figure 4**A** sketches this resolution procedure. In this simple example, the resolving procedure allows one to recover perfect binary trees. However, it is important to note that many incompatibilities due to polytomies are not found in this way. This is especially true when reassortment and large polytomies are entangled in the same clades. Overall, poor resolution represents a loss of information regarding the genealogies, and makes the problem of finding reassortments intrinsically harder.

## Code availability

The code of TreeKnit is available at https://github.com/PierreBarrat/TreeKnit. It is implemented in Julia, and also provides a simple CLI script that returns an ARG file as an extended Newick string [18]. Other codes that were used for this work are listed here:

- The implementation for the simulation of ARGs can be found at https://github.com/PierreBarrat/ARGTools
- Miscellaneous functions that were used to evaluate the performance of TreeKnit are available at https://github.com/PierreBarrat/TestRecombTools.

## III. Results

### Validation on simulated genealogies

We simulated two segment ARGs of one hundred leaves using a coalescent-reassortment process for different values of a scaled reassortment rate *r*^*\**^: coalescence dominates reassortment when *r*^*\**^ ≪ 1, and reassortment dominates coalescence when *r*^*\**^ ≫ 1. Details of the simulations can be found in the SI. Individual segment trees are then extracted from simulated ARGs, and we use TreeKnit to infer MCCs and compare them to the ground truth.

Figure 3**A** shows the number of MCCs *N*_*mcc*_, both real and inferred, as a function of *r*^*\**^. Note that by definition, each MCC corresponds to one reassortment in the ARG, except if it contains the root of both trees. Therefore, the number of reassortments is equal to either *N*_*mcc*_ or *N*_*mcc*_ − 1. As expected, the real number of MCCs (marked black line on the figure) is 1 for very low reassortment rates, and equal to the number of leaves for very high reassortment rates (see Figure 1 for an example).

**Figure 3.**
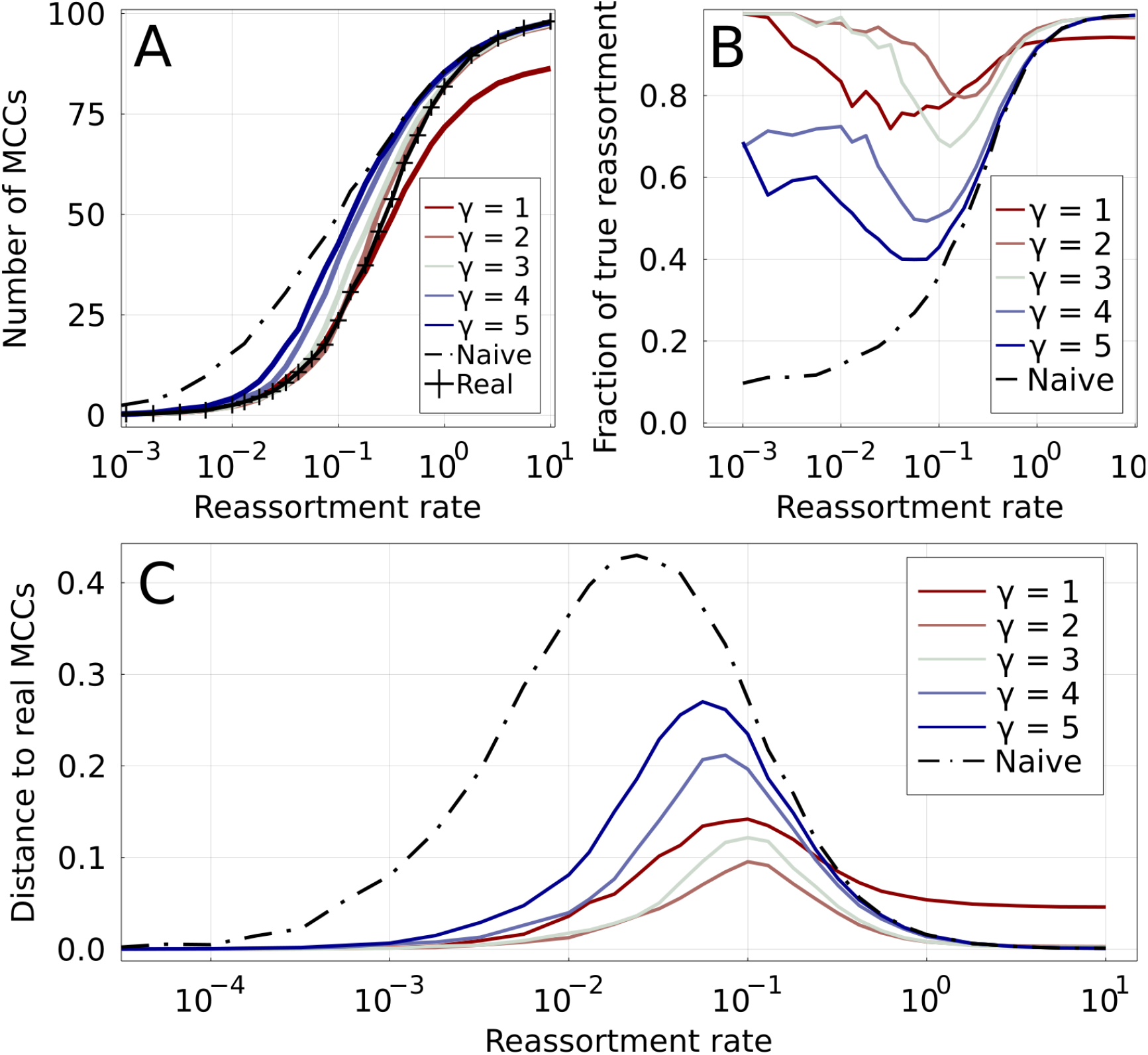
Accuracy of inferred MCCs in simulated ARGs. Increasing values of *γ* are shown by colored lines, from red to blue. **A**: Number of MCCs found by different methods as a function of the reassortment rate. The real number of MCCs is represented by the marked black line. The naive method (dashed black line) overestimates the number of MCCs for low *r*^*\**^, while the parsimonious one (*γ* = 1) underestimates it for high *r*^*\**^. **B**: True positive rate for reassortments: fraction of inferred reassortments that are indeed present in the real ARG. The low number of reassortments results in a relatively large uncertainty for this quantity for *r*^*\**^ ≪ 1. **C**: Distance between inferred and real MCCs for different methods. The distance is based on the variation of information [19].

At fixed *r*^*\**^, the number of inferred MCCs varies systematically with the parameter *γ*. The naive method (*γ* → ∞) is by construction conservative when merging MCCs and consistently overestimates the number of MCCs for *r*^*\**^ *≲* 1 (see discussion in Section II). In contrast, the parsimonious method (*γ* = 1) estimates the number of reassortments for low *r*^*\**^ accurately, but clearly underestimates it for *r*^*\**^ *≳* 0.5. Intermediate values of *γ* fall between these two extremes, with *γ* = 2 and *γ* = 3 being particularly close to ground truth.

Inferring the correct number of MCCs does not necessarily imply that they are correct. Figure 3 shows the positive predictive value (PPV), the fraction of correctly predicted reassortments, as a function of *r*^*\**^. For *γ* equal 2 or 3, the three regimes discussed on Figure 1 are immediately visible: inference is trivially easy in the regions of high and low *r*^*\**^, and the PPV is then close to one. For *γ* = 1, the PPV plateaus below one for high *r*^*\**^ meaning that TreeKnit not only infers to few reassortments, but some of those inferred are incorrect. ARG inference is hardest at intermediate reassortment rates. But even in this region the PPV is above 70% for *γ* = 1 and *γ* = 3, and above 80% for *γ* = 2. This suggests that reassortments are predicted with good accuracy by our method if *γ* is chosen in the right range of values. High values of *γ* overestimate the number of reassortment events at low *r*^*\**^, resulting in a low *PPV*.

The fraction of correctly predicted reassortments is only a limited measure of how well the ARG is recovered as all events, whether deep in the tree or on a terminal branch, are weighted equally. We propose an alternative measure to globally assess the accuracy at which the inference represents the truth based on the idea that MCCs define a *clustering* of leaves. Two clusterings of the same set can be compared using the variation of information (VI) [19]. The VI of two partitions of a set is equal to the difference between the sum of the entropies of the two partitions and their mutual information times two. It is 0 if the two partitions are identical, and equal to log(*L*) if the two partitions are maximally different, where *L* is the number of elements of the set, here the number of leaves in the trees. Here, we scale the VI by log(*L*) so that it varies between 0 and 1. While the values of VI are not as directly interpreted as a fraction of true positives, they constitute a global measure of how close a set of inferred MCCs is to the ground truth and can thus be used to compare methods.

Figure 3**C** shows the VI distance between the real MCCs and the ones inferred by the TreeKnit for different *γ*. Regions where *r*^*\**^ ≫ 1 and *r*^*\**^ ≪ 1 result in accurate inference for *γ >* 1. As expected, the performance of the naive method starts to decrease as soon as *r*^*\**^ rises above very small values, or in other words as soon as reassortments appear in the ARG. As before, *γ* = 2 seems to be an optimal value across the entire range of reassortment rate. In the SI, we show that *γ* = 2 to 3 is optimal for different coalescent processes (Figures S2 & S3). Therefore, fine-tuning of *γ* does not seem necessary.

Another way of estimating the accuracy of the inference is to look at predictions made for individual branches of the segment trees. For one of the trees, each branch can either be predicted to be shared with the other tree in the ARG, or to be specific to its tree (see illustration in Figure 1). Figure S1 in the SI shows the accuracy at which individual branches are predicted to be shared or not for different values of *γ*. As expected, *γ* = 1 results in branches incorrectly predicted to be shared, while for large values of *γ* shared branches are often missed. The values of *γ* lying in between seem to smoothly interpolate between these two behaviors. *γ* can thus be seen as a way to balance between different types of false predictions for branches of the trees.

### Increasing phylogenetic resolution of segment trees

While simulated trees are completely binary, trees reconstructed from genome sequences often lack resolution resulting in polytomies. This problem is particularly acute when the sample consists of many closely related viruses. As stated in section II, poor resolution of trees leads to complications for our topology based method: topological differences due to reassortment must be disentangled from those due to different polytomies in segment trees, see Figure 4**A** for example. Polytomies are thus a source of errors in the inference of MCCs. At the same time, the knowledge of the MCCs allows us to better resolve trees: in regions of shared branches in the ARG, the two trees must have the same splits, *i*.*e*. the same internal nodes. This means that once the MCCs are inferred, it is possible to introduce some of the splits of each tree in the other, and therefore to partly resolve them.

To quantify the errors in ARG reconstruction due to limited resolution and to assess the quality of the resolution once the MCCs are known, we simulated ARGs with limited resolution (see A 4 of the SI). A parameter *c* ≥ 0 controls the amount of polytomies: *c* plays the role of an inverse mutations rate and *c* = 0 corresponds to perfectly resolved binary trees where every branch is supported by mutations, while *c* → ∞ corresponds to completely unresolved star-trees. A value of *c*^*\**^ ≃ 0.8 results in a level of resolution that is quantitatively close to the one observed in trees of A/H3N2 HA with strains of the same season (see Figures S5 and S6).

Figure 4**B** shows the VI distance of inferred MCCs to the real ones (using *γ* = 2) as a function of *r*^*\**^ for varying resolution of the segment trees controlled by *c*. The case of *c* = 0 is equivalent to the *γ* = 2 curve on Figure 3**C**. The performance of the method decreases gradually with the loss of resolution. For A/H3N2 like resolution (*c* = 0.8), the error is close to doubled when compared to the *c* = 0 case. However, for most reassortment rates, the gain in performance over using the naive method remains substantial.

Once MCCs are inferred, we can resolve the subtrees corresponding to an MCC by complementing each others splits. However, this will only result in correct resolutions if the MCCs are correctly identified to begin with and thus requires further validation. Figure 4**C** shows the number of missing splits that are correctly and incorrectly introduced scaled by the total number of missing splits, as a function of *r*^*\**^ (lines with markers) and for both the naive and the *γ* = 2 method. Note, that a split that is resolved in neither tree can not be resolved this way. The maximal resolution, obtained with the knowledge of the true MCCs, is indicated in the figure, showing that for *γ* = 2 the majority of possible resolutions are found while only a small fraction of incorrect splits are introduced.

### Comparison with other methods

To demonstrate the utility of TreeKnit, we compare it to GiRaF [13, 20] and the recently published fully bayesian method CoalRe [8, 16].

GiRaf is, as TreeKnit, based on topological differences between the two trees. Given two trees, it uses the compatibility network of their splits to infer the position of reassortments. However, unlike our method, it does not aim at inferring the whole ARG. Instead, it finds a set of probable reassortments, which may not fully explain the topological differences between segment trees.

CoalRe [8, 16], on the other hand, is a bayesian method that uses a coalescence-reassortment process to construct a probability distribution for ARGs given a sample of sequences. It reconstructs the ARG by sampling from this distribution. It can thus be seen as an extension of usual tree inference methods to the case of genealogies with reassortment. By construction, it uses not only the topology of segment trees, but of all the available information in the form of gene sequences.

We compare methods for three different reassortment rates: *r*^*\**^ ∈ {0.01, 0.05, 0.25}. Note that the intermediate value *r*^*\**^ = 0.05 is close to the one that we estimated for segments HA and NA of A/H3N2 influenza. For each value of *r*^*\**^, we simulate five ARGs and extract the segment trees from them. We then simulate the evolution of sequences on these trees, using the JC69 model for simplicity. Mutation rate *μ* is either tuned such that the resulting trees have a close to perfect resolution (*μ*^*high*^), or such that the trees have flu-like polytomies (*μ*^*low*^). We use all three methods to infer reassortment from the simulated sequences. In the case of GiRaF and TreeKnit, trees must first be reconstructed: this is done using the MrBayes program [21] in the case of GiRaF (code provided in the original publication [13]), and using IQ-Tree in the case of TreeKnit [22]. The natural inputs of CoalRe are sequence alignments, and no prior tree reconstruction is needed.

Figure 5 quantifies the accuracy and completeness of the inferences made by the tree methods. TreeKnit and CoalRe perform similarly well, with CoalRe reporting slightly less false reassortments. GiRaF misses reassortment events, but very rarely reports a reassortment that did not happen.

**Figure 5.**
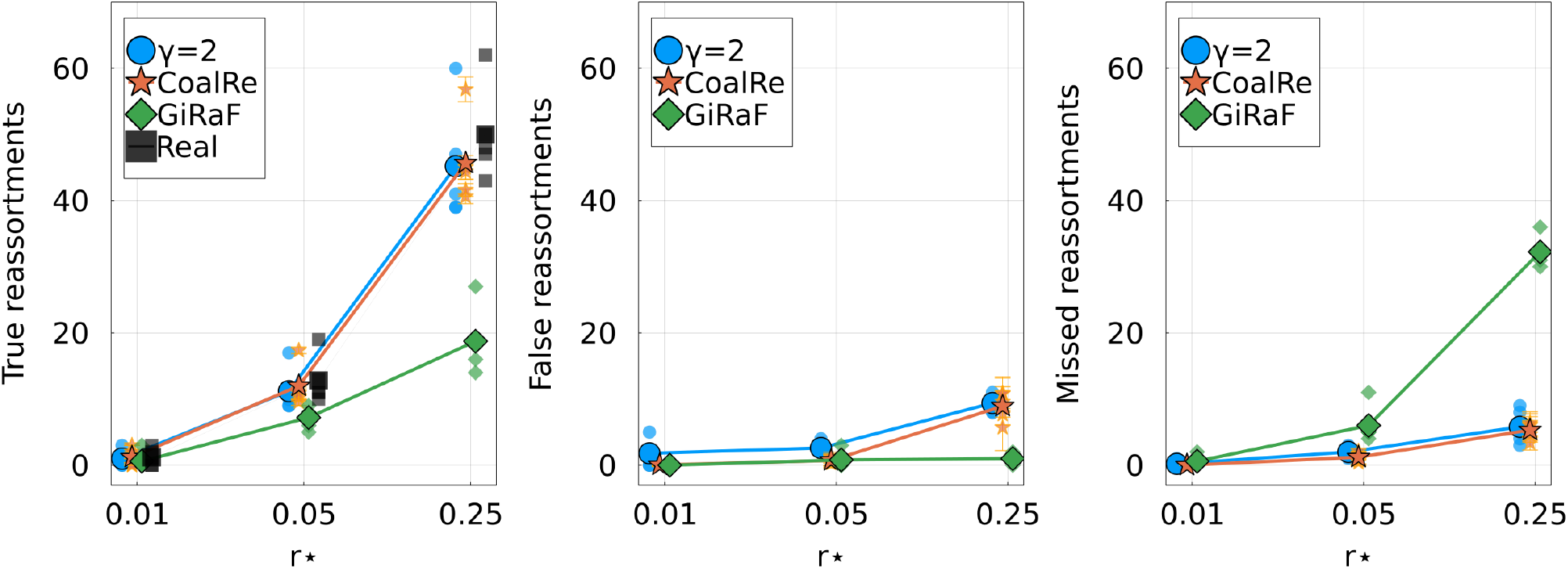
Comparison of TreeKnit with CoalRe [8, 16] and GiRaF [13, 20] on simulated ARGs of 100 leaves. For three reassortment rates, shows the **Left**: number of true reassortments, **Center**: number of false reassortments, and **Right**: number of missed reassortments, for all three methods. Large markers represent the average over 5 simulations for each *r*^*\**^. Smaller markers show results on individual ARGs. Results for each method are slightly shifted on the *r*^*\**^-axis for visibility.

TreeKnit has a much shorter runtime than the other methods. For the trees of 100 leaves used here, the average runtime of TreeKnit was 40ms, whereas GiRaF took 40s. Note that these times are small compared what is needed to infer the phylogenetic trees: the IQ-Tree runs took about 30s, while the MrBayes runs took about 20minutes. Figure S10 shows that the runtime of TreeKnit scales quadratically with the number of leaves. We also observed that GiRaF scales similarly. Finally, CoalRe takes several hours to infer one ARG. TreeKnit thus runs orders of magnitude faster than CoalRe at the cost of a very small reduction accuracy.

### Results on influenza A data

TreeKnit was developed for application to influenza virus data and in particular to seasonal influenza viruses. We have shown (see Figure 3) that TreeKnit accurately infers MCCs of simulated genealogies. But accuracy depends on the reassortment rate and the resolution of the segment trees. In the SI, we estimate that A/H3N2 influenza data corresponds to *r*^*\**^ ≈ 0.05 and *c* = 0.8 (Figures S4**A** S5). Figure 4**B** uses simulations to estimate the difficulty of the inference problem in the space of *r* and *c*. Seasonal influenza viruses seem to lie exactly in a region of parameter space where the problem is hard, but most is gained over the naive method. We can thus expect significant improvement in terms of resolution of the segment trees and non-trivial reassortment patterns.

We illustrate the use of TreeKnit on the collection of human influenza H3N2 isolates sequenced and analyzed in [9]. The data consists of about 150 strains sampled in New York between 1999 and 2004. This data has previously been used as a test set by some authors [7, 13].

When considering HA and NA segments, two reassorted clades were found in [9] to be reassortments by manual inspection. The GiRaF method [13] found more reassorted clade containing a single strain. We applied TreeKnit to this dataset (*γ* = 2), with results shown on Figures S8 and S9. TreeKnit recovers the two reassortments found in [9] and the additional one found by GiRaF, and also finds three extra reassorted clades: {A/New York/137/2004, A/New York/138/2003}, {A/New York/177/1999}, and a large clade consisting of strains sampled between 2003 and 2004. Manual inspection of the trees suggests that these three clades are indeed reassortments. Note that the fact that we find more reassortments than GiRaF on the same data is consistent with results shown on Figure 5.

## IV. Discussion

Recombination and reassortment are important to reduce mutational load by weeding out deleterious mutations [23] and can facilitate adaptation, especially in rapidly evolving popululations [24, 25]. While a number of studies document frequent reassortment in seasonal influenza virus populations [6, 7, 9, 26], reassortment is often ignored because of a lack of methods to infer ARGs at scale. Furthermore, the inferred reassortment history is difficult to visualize and interpret as soon as more than a few reassortment events occurred.

We introduced TreeKnit, a fast and accurate method to infer ancestral reassortment graphs (ARG) from two segment trees. Its underlying idea is to find regions of the genealogy without reassortments, where the two trees have the same topology (MCCs). Intuitively, this is done by “knitting” the trees together starting from the leaves. The method iteratively grows compatible regions and introduces reassortment events until the entire observable ARG is inferred. At each stage, the method introduces reassortment events that explain the largest number of incompatibilities and stops when no more reassortment events are found that explain at least *γ* incompatibilities.

The parameter *γ* allows us to tune the behavior of TreeKnit: at *γ* = 1, TreeKnit tries to minimize the number of reassortments, while at *γ* → ∞ only complete subtrees with identical topology are identified. Simulations revealed that *γ* = 2 is a robust choice, giving best results a wide range of reassortment rates. Note that the method is by design *greedy*, since the function 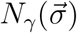 only takes into account incompatibilities that are one level above the (effective) leaves. Comparing TreeKnit to CoalRe suggests that the greedy heuristic achieves similar accuracy to more principled bayesian approaches for parameter sets tested, while being orders of magnitude faster.

Efficient inference of ARGs of seasonal influenza viruses should enable deeper insights into the importance of reassortment for antigenic evolution through combination of specific HA and NA variants. So far this has remained unclear, with studies reaching different conclusions [7, 8]. This discordance might in part due to the lack tools that can infer ARGs of large data sets. In addition, joint analysis of multiple segments should improve resolution of segment trees with polytomies. In parts of the ARG that are common to both segment trees, the sequences of the two segments can be considered jointly, which amounts to roughly doubling the number of mutations per branch. In the future, we plan to extend TreeKnit to multiple and complement it with ARG visualization tools in Nextstrain [27].

## A. Detailed methods

### 1. Minimization of *N*_*γ*_ using simulated annealing

Minimizing 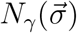 amounts to a discrete optimization problem. The functional form of *N*_*γ*_ in Equation (1) allows us to quickly compute a value for a given configuration 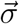, but does not seem to be amenable to simple optimization. In particular, it is not clear whether many local minima exist. For this reason, we rely on the general technique of simulated annealing [17].

The algorithm consists having configurations 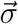 perform a random walk in the energy landscape defined by 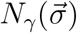 at a “temperature” *T*, using a simple Markov Chain Monte Carlo (MCMC) method. This is equivalent to sampling from the probability distribution 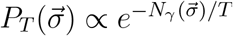. The temperature *T* is initialized at a high value and slowly brought to 0: this is the cooling process. For an infinitely slow cooling, the sampling process will converge to 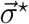, the minimum of *N*_*γ*_.

#### Convergence and reproducibility

In practice, cooling cannot be infinitely slow, and there is no guarantee that the algorithm converges to the global minimum, or even that two subsequent runs converge to the same point. However, we observe in our setting that annealing runs give highly reproducible results as long as the cooling speed is inversely proportional to the number of leaves in the trees, that is if the number of iterations of the SA algorithm is proportional to the number of leaves *L*. Figure S11 shows the distance of inferred MCCs to the real ones as a function of the number of iterations at each temperature *T*, scaled by the number of leaves *L* in the ARG. It is clear that the optimization converges for a high enough number of iterations, and that the required number of iterations for proper convergence should be proportional to *L*.

Reproducibility of results is shown on Figure S12: the distance between MCCs obtained in two independent runs on the same data is plotted against the number of iterations performed at each temperature *T*, scaled by the number of leaves *L*. Since the starting point of the optimization is the result of the naive method, the difference between two runs is low for a low number of iterations. It goes through a maximum for an intermediate number, and vanishes again as the optimization converges. The fact that all curves “stack” vertically shows that the number of iterations of the SA algorithm should be chosen proportionally to *L*.

#### Runtime

The computational complexity is of the minimization can easily be estimated. Computing *N*_*γ*_ for a tree of *L* leaves results in order *L* operations. Since we choose the cooling speed of the SA to be inversely proportional to *L*, we obtain a quadratic complexity 𝒪 (*L*^2^). The minimization of *N*_*γ*_ is the most computationally intensive part of the algorithm for large trees (*L ≳* 100). As a result, we expect the overall runtime to be quadratic in *L*. This is verified in Figure 10.

### 2. Simulation of ARGs

We simulate ARGs using a backwards coalescence-reassortment process. The process is initiated with *n*_0_ leaf nodes that all have two segments. Two types of event can then occur:

- Coalescence, with a rate *ν*_*c*_(*n*), where *n* is the number of remaining lineages in the simulation (*n*_0_ at start). In a coalescence, two nodes are picked at random and their lineages are merged. If these lineages corresponded to different segments or if at least one of the lineages corresponds to two segments, the ancestral node formed will have two segments. Otherwise, it has one. The choice of a functional form for *ν*_*c*_(*n*) is detailed below.
- Reassortment, with a rate *ν*_*r*_ = *rn*_*r*_ with *r* constant. *n*_*r*_ is the number of nodes that have two segments. In a reassortment, a node with two segments is chosen at random and its lineage splits backward, giving rise to two ancestors. Each segment goes to one of these ancestors, and these can no longer be reassorted as they only have one segment.

Sampling of events continues until a root for each segment has been found.

The *scaled reassortment rate* is defined using the ratio of reassortment to coalescence rates at the start of the simulation:

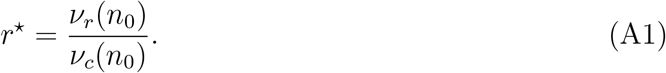

Importantly, *r*^*\**^ describes the competition between coalescence and reassortment at the very start of the simulation, when all leaves are present. Consequently, the value *r*^*\**^ = 1 means that the *first event* that will be simulated is equally likely to be a coalescence or a reassortment.

The rate of coalescence can be chosen in several manners. The Kingman coalescent is defined by *ν*_*c*_(*n*) = *n*(*n* − 1)*/*2*N*, with timescale *N* corresponding to a global population size. It is characterized by very short branches close to the tips of the ARG, and long branches that lead back to the root. The Yule coalescent is defined by *ν*_*c*_(*n*) = (*n* − 1)*/N*, and typically has much longer terminal branches than the Kingman’s. For results of the main text, we use a custom coalescent model that is designed to reproduce the distribution of tree branch length that is empirically observed in A/H3N2 influenza genealogies. It is defined by *ν*_*c*_ = *n*^0.2^(*n* − 1)*/*2*N*. All results presented in this article are qualitatively unchanged when using these different types of coalescent models. Figures S2 and S2 are equivalent to Figure 3 from the main text, and show results obtained using the Yule and the Kingman coalescent models.

### 3. On the reassortment rate

We explain here the choice of the definition of *r*^*\**^ in Equation A1, and the difference with the usual definition of the reassortment rate *ρ* in literature. In the literature, the reassortment rate *ρ* is usually defined as *rT*_2_, where *T*_2_ is the pairwise coalescence time (with our notation, we have *T*_2_ = *ν*_*c*_(2)^−1^). In other words, *ρ* is the average number of reassortments occurring on a lineage during the time it takes for a pair of leaves to coalesce to a common ancestor. If the value of *T*_2_ is known in years, as in the case for the influenza virus, then *ρ* is measured in reassortments per lineage per year.

However, using *ρ* as a control parameter for the strength of reassortment in simulations is impractical: at a fixed *ρ*, the properties of the MCCs will vary with *n* and with *ν*_*c*_. As an example, consider an ARG generated using a Kingman coalescent: *ν*_*c*_(*n*) = *n*(*n* − 1)*/N*. Adding more leaves to the simulation will result in an initial coalescence rate that is much higher than the reassortment rate, and many coalescences will take place before the first reassortment. As a result, going from a low *n* to a large *n* will take us from a situation like the one in the right panel of Figure 1 (many reassortments, small MCCs, very different segment trees) to one like the left panel of the same figure (very few reassortments, large MCCs, very similar segment trees). *ρ* alone is therefore not a good indicator of the different regimes shown in Figure 1.

For this reason, we construct another reassortment rate *r*^*\**^, defined in Equation A1, which controls the relative strength of coalescence and reassortment. *r*^*\**^ is indicative of the different regimes depicted in Figure 1: the transitions between low, intermediate and high levels of reassortments will take place in the region *r*^*\**^ ≃ 10^−2^ − 10^−1^, irrespective of the number of leaves and of the coalescent model used. In particular, the choice of *r*^*\**^ as a scale for the reassortment rate allows Figures 3, S2 and S3 to use the same range of values on the *x*-axis, even though they correspond to three different coalescent models.

*ρ* and *r*^*\**^ are related in a straightforward way. First, we write the coalescent rate as follows:

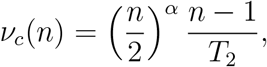

with the Kingman coalescent corresponding to *α* = 1, the Yule coalescent to *α* = 0, and our custom coalescent model to *α* = 0.2 (up to a proportionality constant). Note that *T*_2_ defines the pairwise coalescence time *T*_2_ = *ν*_*c*_(2). From there, we immediately obtain

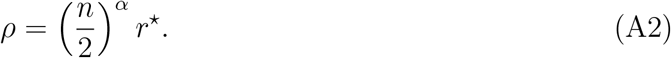

In Figure 4**A**, we estimate *r*^*\**^ ≃ 0.06 in the case of segments HA and NA of A/H3N2 influenza. Since this was estimated using a coalescent with *α* = 0.2 and with *n* = 100 leaves, we have for the reassortment rate *ρ* ≃ 0.13 reassortments per lineage per year. This is smaller but in the same order of magnitude than estimations obtained by the CoalRe method [8].

### 4. Introducing polytomies in simulated trees

To investigate the case of incompletely resolved trees, we need to introduce polytomies in the trees that come from the simulated ARG. We do so in the following way: for each each branch of length *t* in a given tree, we remove it with an exponentially decreasing probability *P*_*r*_ = exp (−*t/cN*). Here, *c* is a parameter that can be varied between 0 (no branches removed, binary trees) and ∞ (star trees, completely unresolved), and *N* is the population size that was used in the coalescent model. If sequences were simulated on the trees by a simple evolutionary process, this would amount to remove branches with no mutations, with *cN* acting as the inverse of a mutation rate.

Figure S5 shows the ratio of the number of leaves to the number of internal nodes for A/H3N2 HA trees (dashed lines) as well as for simulated ones with varying values of *c* (solid lines). This ratio should be one half in the case of perfectly resolved trees, which are then binary, and can go up to one for fully unresolved trees (star-tree). As expected, the ratio increases with the number of strains used to build the trees. The figure shows that a unique value *c*^*\**^ allows us to replicate the ratio of leaves to internal nodes of HA trees for different values of the number of leaves.

Note that this result is a consequence of the specific coalescent-reassortment model that we used, discussed in section SA 2. In particular, a different choice of the coalescence rate would make *c*^*\**^ depend on the number of leaves. In addition, Figure S6 shows that the simulated trees can closely reproduce the distribution of polytomy sizes. Together, these results show that we can reliably replicate the lack of resolution of influenza gene trees in our simulations, and therefore estimate its effect on the reconstruction of the ARG.

### 5. Choosing among equivalent topological solutions using a likelihood test

Our algorithm, schematized in Figure 2, consists in optimizing the discrete function *N*_*γ*_ as a function of the set 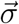 of effective leaves removed or kept in the tree. The optimal configuration 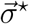 gives us the leaves to remove, that is the found MCCs for this iteration of the algorithm, and the leaves to keep. For certain input trees or reduced trees, it is possible that *N*_*γ*_ has several minima. These corresponds to sets of putative MCCs that are equivalently good solutions in terms of topology only. A very simple case of such a situation is for the set of the two following trees (written as Newick strings, without branch length for simplicity): `((A,B),C)` and `(A,(B,C))`. In this case, three incompatibilities exist when no leaf is removed, and removing any of the leaves `A`, `B` or `C` fixes all incompatibilities. Therefore, for *γ <* 3, three topologically equivalent solutions are found by the algorithm.

In such cases, we need to use information beyond topology to break the degeneracy of solutions. The most intuitive information at our disposal is of course branch length. In order to use branch lengths to pick one solution, we derive here a simple likelihood test. First, we make some simple hypothesis over the evolutionary process that gave rise to the tree and over the signification of branch lengths. For each segment *i*, we call *L*_*i*_ the length of its sequence, *μ*_*i*_ its per site mutation rate, {*t*_*i*_} the length of branches of its tree. The branch length *t*_*i*_ is interpreted as the number of mutations *per sequence site* that occurred on said branch.

For a given branch in the tree of segment *i*, we call *T*_*i*_ the physical time corresponding to this branch. By definition of the mutation rate, the expected length of this branch in the tree is therefore ⟨*t*_*i*_⟩ = *μ*_*i*_*T*_*i*_. However, the actual observed value of *t*_*i*_ may not be equal to the expected one because of the stochastic nature of mutation events. We assume here that the probability of observing a given number of mutations *n*_*i*_ = *L*_*i*_*t*_*i*_ on this branch is given by a Poisson distribution with mean 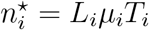:

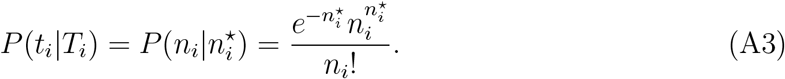

We now make two observations about our algorithm. The first is that removing a leaf from a configuration 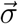 amounts to making that leaf an MCC and introducing a reassortment above it. Therefore, the associated prediction is that the branches joining said leaf to its ancestors in the two segment trees are not shared in the ARG and can be of two different lengths and correspond to two different physical times. We can attribute a probability of observing branch lengths *t*_1_ and *t*_2_ above said leaf given that we predict these branches are not shared:

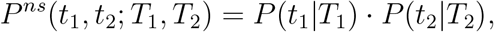

where *T*_1_ and *T*_2_ are the physical times corresponding to the two branches. The most likely values of *T*_1_ and *T*_2_ are those that respectively maximize *P* (*t*_1_|*T*_1_) and *P* (*t*_2_|*T*_2_), giving 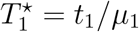 and 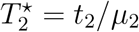. We therefore define the likelihood of the “not shared branches” prediction as

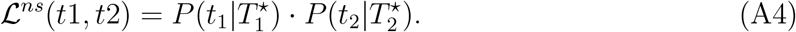

The second observation is that if there is no incompatibility for a given leaf, the prediction of the algorithm is that no reassortment occurred above it. Therefore, the branches above the leaf will be shared in the ARG, and must correspond to the same physical time *T*. Proceeding as above, we compute the probability of observing branch length *t*_1_ and *t*_2_ if predicting that they are shared branches:

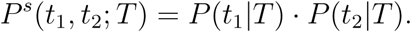

It can easily be shown that if the mutation rates are similar, the most likely value of *T*, that is the one maximizing *P* ^*s*^(*t*_1_, *t*_2_; *T*), is

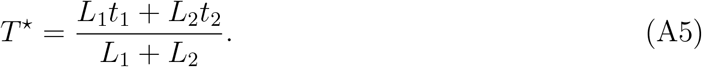

In turn, this allows us to define a likelihood for the “shared branches” prediction:

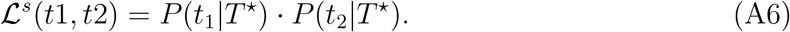

For a given configuration 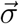, we now proceed as follows to score it using branch lengths. For each leaf *n* such that *σ*_*n*_ = 0, we count a likelihood ratio score of *L*^*ns*^(*t*_1_, *t*_2_)*/L*^*s*^(*t*_1_, *t*_2_), where *t*_1_ and *t*_2_ are the branch lengths above *n* in each segment tree. Inversely, for each leaf *n* such that *σ*_*n*_ = 1 and that does not have any incompatibility above it, we count a likelihood ratio score of *L*^*s*^(*t*_1_, *t*_2_)*/L*^*ns*^(*t*_1_, *t*_2_). These scores are multiplied, giving us an overall score for 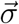. When several optimal configurations are found after optimization, the one with the highest likelihood ratio score is chosen.

Even though this likelihood test is necessary to the completeness of our method, we find in practice that it has little influence over the results. The quality of the inferred MCCs is only marginally improved when using it. This likely comes from the relatively infrequent occurrence of conflicting degenerate minima of *N*_*γ*_.

### 6. Simulating sequences

For comparison with CoalRe. Nucleotide sequences of length 1000 are simulated independently on each segment tree, using a JC69 model.

### 7. Robustness with regard to tree inference

The method described in this article relies entirely on topological differences between trees. This means that topological mistakes done in the reconstruction of the trees from sequences can have an important impact on the results. In particular, branches that are not well supported will tend to introduce many spurious incompatibilities between trees.

To mitigate this effect, we pre-process trees obtained from A/H3N2 influenza sequences in the following way:

- If sequences are of length *L*, every branch shorter than 1*/*(2*L*) are removed. This strongly reduces the number of branches on which a mutation is unlikely.
- Branches with a bootstrap support smaller than *b*_0_ = 75 are removed (ultra fast bootstrap test in iqtree [4]).

To determine *b*_0_, we used the following test. For a given alignment, we infer two trees independently, keeping only branches with bootstrap support higher than *b*. We then give these two trees as an input to our method and infer a set of MCCs. In principle, the two trees should be exactly the same, and we should therefore find one MCC containing all the leaves. Figure S13 plots the average number of MCCs found in this way as a function of *b*. It is clear that for *b >* 75, no spurious topological incompatibility remain.

## B. Supplementary figures

**Figure S 1.**
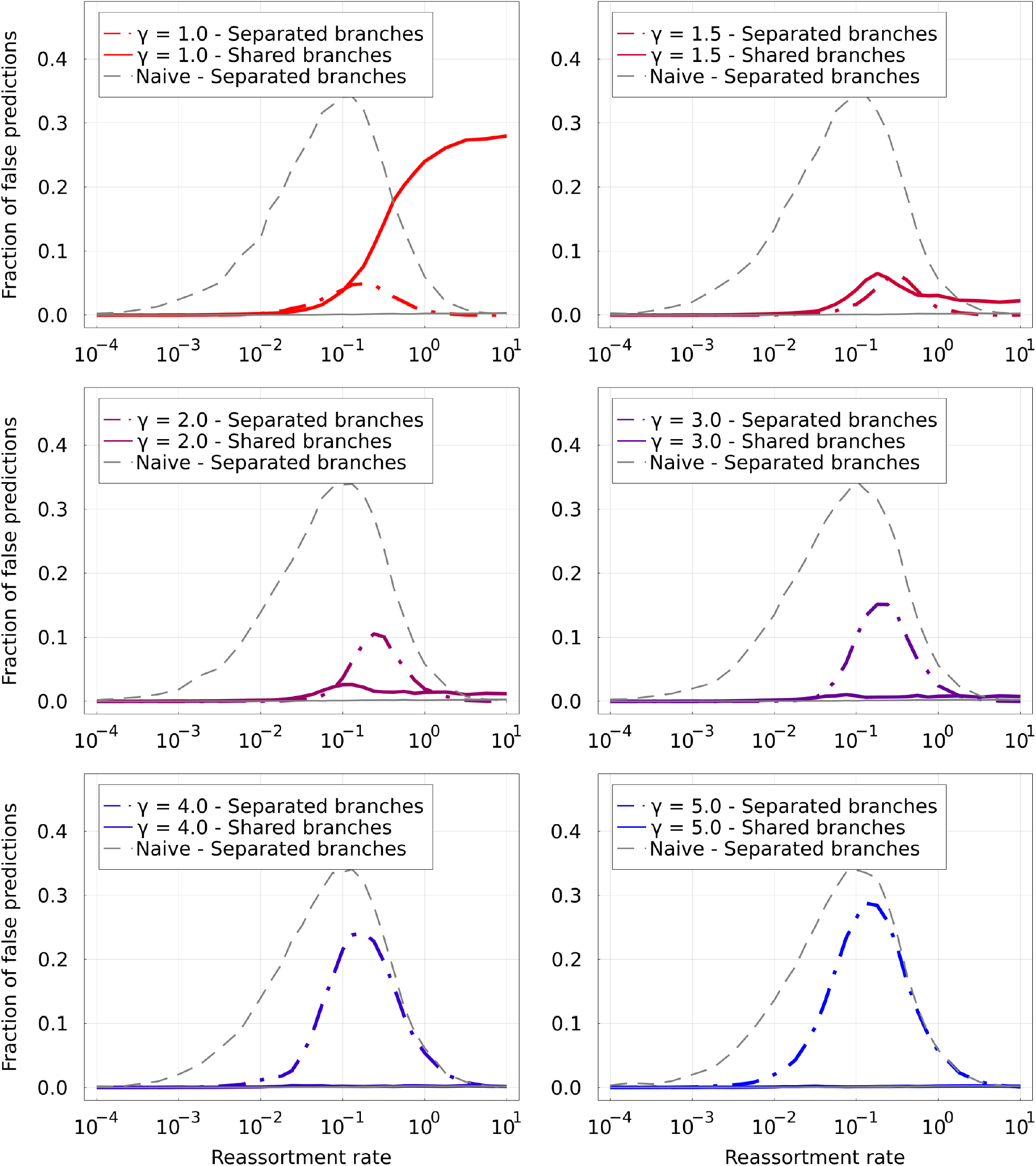
False predictions for individual branches as a function of *ρ*, for different values of *γ*. Solid lines correspond to branches that are predicted to be shared by the two segment trees, but are not in the real ARG. On the contrary, dashed lines correspond to branches that are predicted to not be shared by the two segment trees, but are shared in the real ARG. The parsimonious method (top-right) makes a lot of the first type of error, and few of the second type. Increasing *γ* transitions between the two types of errors. The naive method (grey dashed line) makes close to only errors of the second type.

**Figure S 2.**
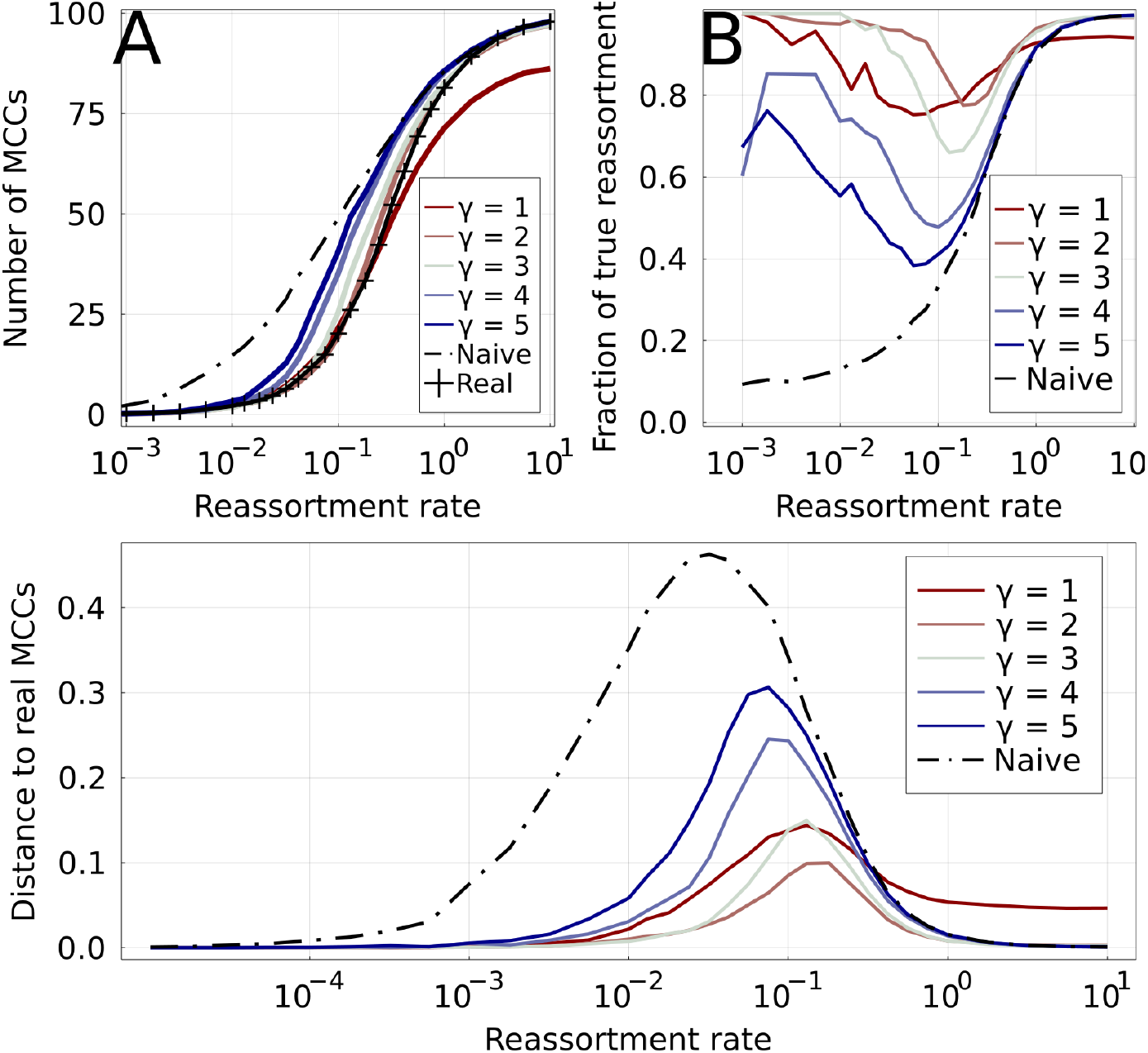
Equivalent to Figure 3 of the main text for the Yule coalescent. Results of *γ*-methods on simulated ARGs of 100 leaves. Increasing values of *γ* are shown by colored lines, from red to blue.**A**: Number of MCCs found by different methods as a function of the reassortment rate. The real number of MCCs is represented by the marked black line. The naive method (dashed black line) overestimates the number of MCCs for low *ρ*, while the parsimonious one (*γ* = 1) underestimates it for high *ρ*. **B**: True positive rate for reassortments: fraction of inferred reassortments that are indeed present in the real ARG. The low number of reassortments results in a relatively large uncertainty for this quantity for *ρ* ≪ 1. **C**: Distance between inferred and real MCCs for different methods. The distance is based on the variation of information

**Figure S 3.**
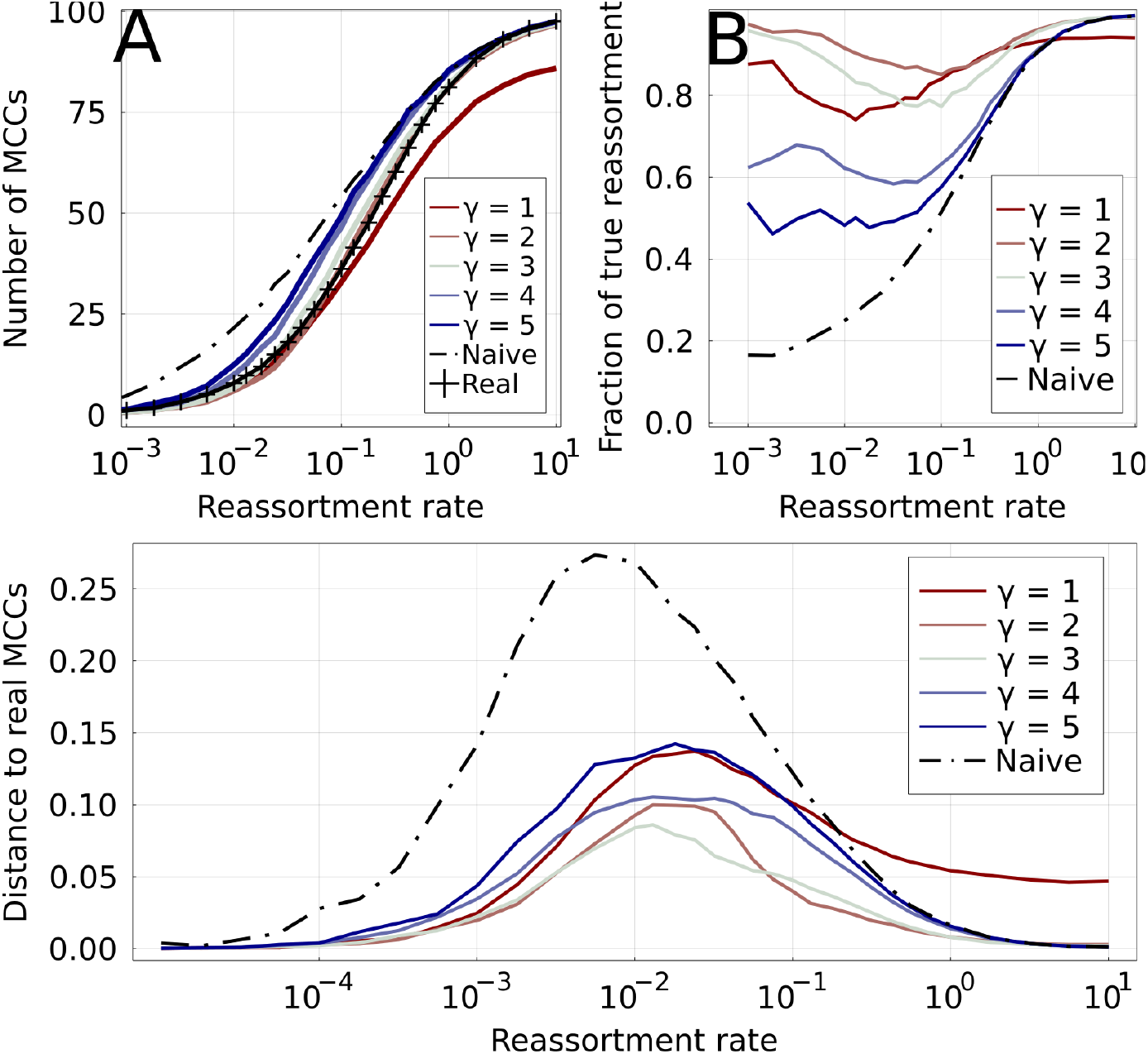
Equivalent to Figure 3 of the main text for the Kingman coalescent. Results of *γ*-methods on simulated ARGs of 100 leaves. Increasing values of *γ* are shown by colored lines, from red to blue.**A**: Number of MCCs found by different methods as a function of the reassortment rate. The real number of MCCs is represented by the marked black line. The naive method (dashed black line) overestimates the number of MCCs for low *ρ*, while the parsimonious one (*γ* = 1) underestimates it for high *ρ*. **B**: True positive rate for reassortments: fraction of inferred reassortments that are indeed present in the real ARG. The low number of reassortments results in a relatively large uncertainty for this quantity for *ρ* ≪ 1. **C**: Distance between inferred and real MCCs for different methods. The distance is based on the variation of information

**Figure S 4.**
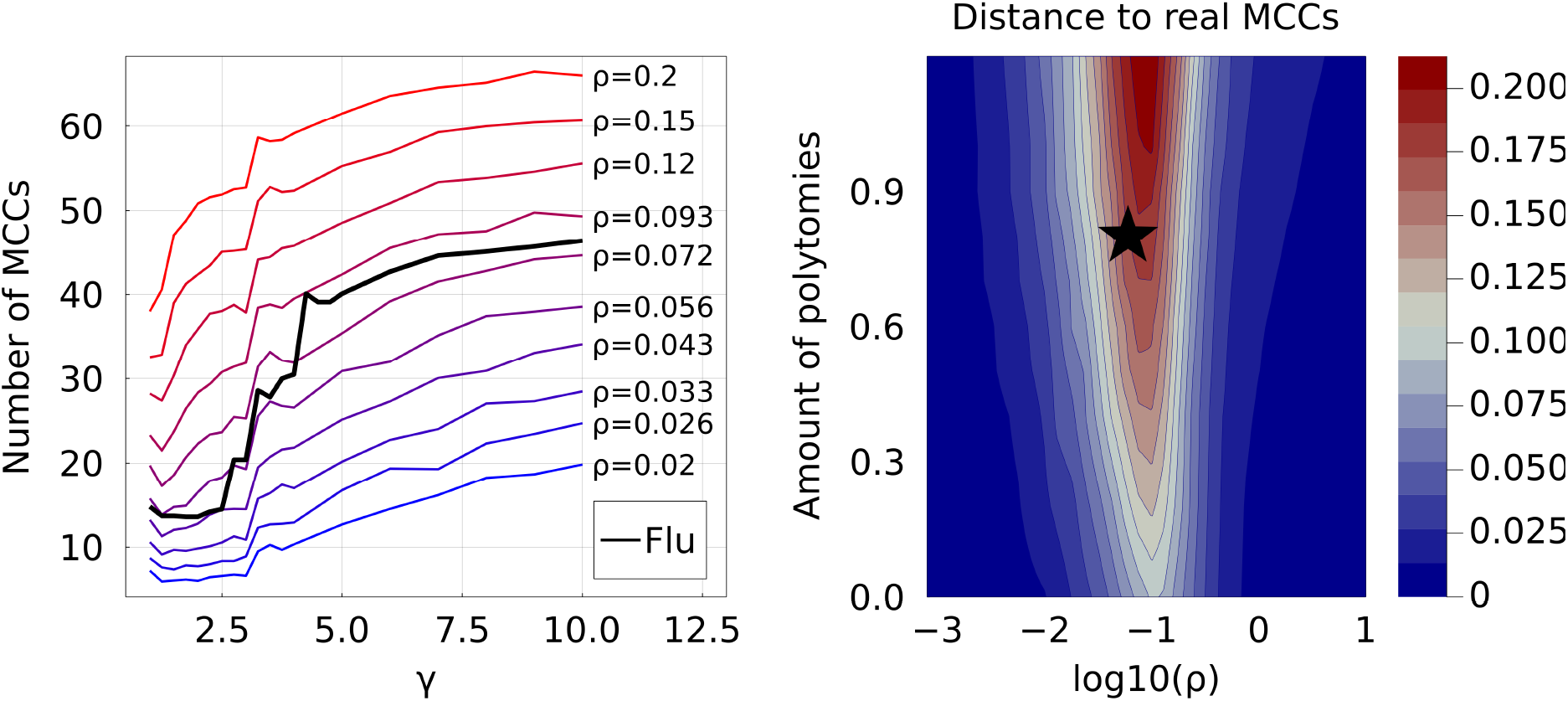
**Left**: Estimation of the reassortment rate of segments HA and NA in A/H3N2 influenza. Colored lines show the number of MCCs inferred by *γ*-methods as a function of *γ*. Colors going from blue to red correspond to increasing values of *ρ*. The black line shows the same quantity for the influenza trees. The curve corresponding to influenza lies between the values 0.043 and 0.093. **Right**: VI distance of inferred MCC to the real ones for simulated data and *γ* = 2, as a function of the reassortment rate *ρ* and the amount of polytomies in the trees. The star corresponds to the estimated position of A/H3N2 influenza (HA/NA segments, sequences from the same epidemiological season).

**Figure S 5.**
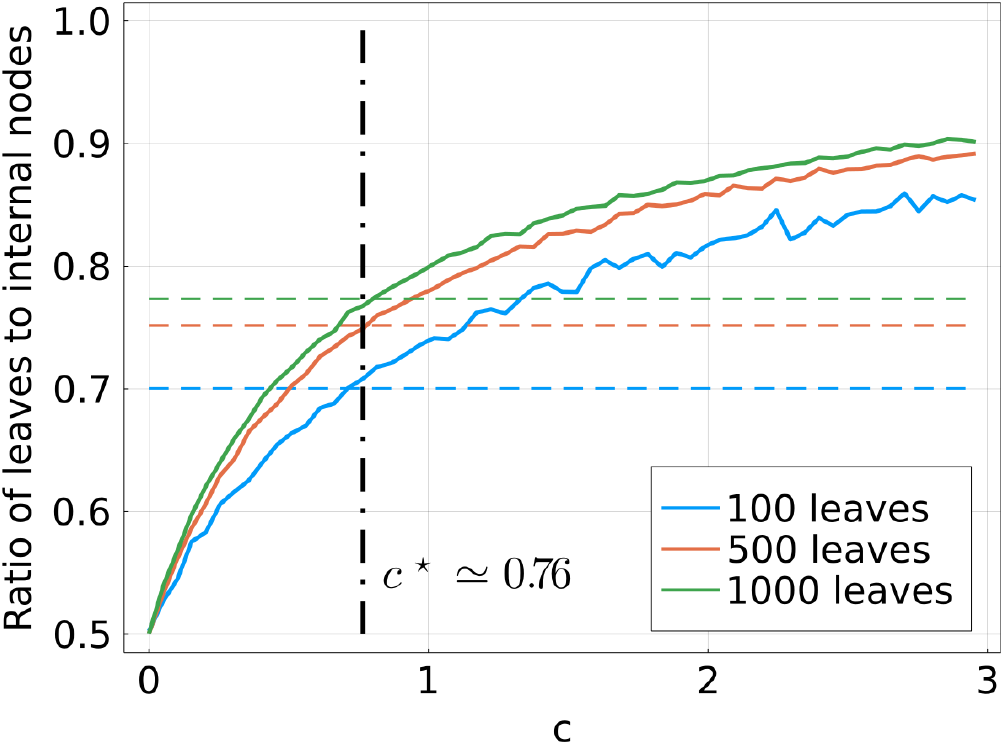
Solid lines: ratio of number of leaves to number of nodes in simulated trees of different sizes, as a function of the amount of parameter *c*, artificially introducing polytomies. Dashed lines: same quantity for A/H3N2 HA trees. For a perfectly resolved tree, this quantity is approximately one half. Polytomies are introduced in simulated trees using the method described in section A 2 of the SI. A unique value *c*^*\**^ allows simulated trees of different sizes to reproduce the lack of resolution of influenza trees.

**Figure S 6.**
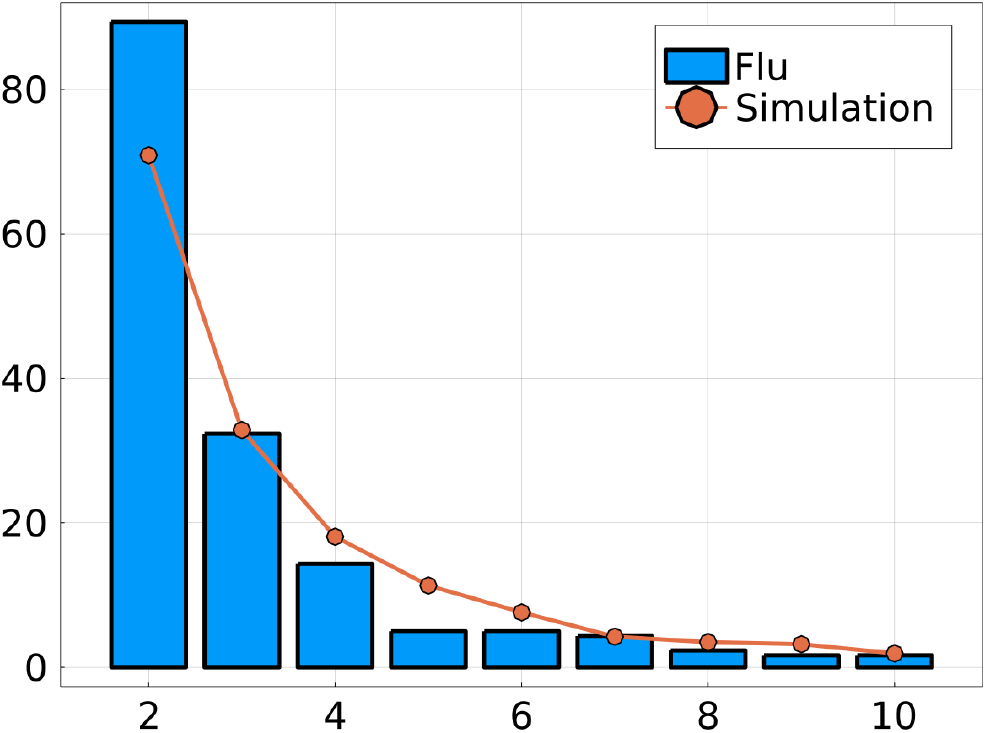
Histogram of the size of polytomies for A/H3N2 HA trees and for simulated ones, with a level of polytomies chosen to reproduce that of influenza.

**Figure S 7.**
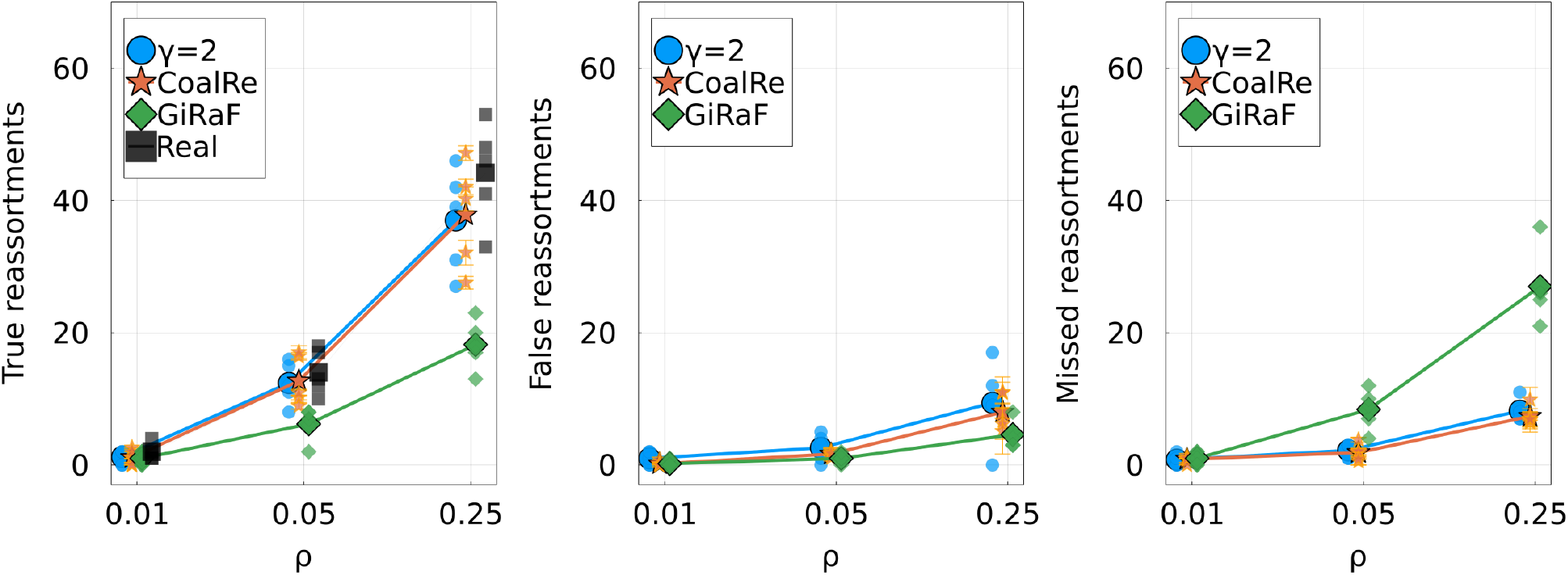
Similar to Figure 5 from the main text, with sequences generated using a lower mutation rate. This results in reconstructed trees with more polytomies. **Left**: number of true reassortments, **Center**: number of false reassortments, and **Right**: number of missed reassortments, for all three methods (Treeknit, GiRaF, CoalRe). Compared to better resolved trees (Figure 5), GiRaF and Coalre infer more false reassortments while Treeknit has more missed reassortments.

**Figure S 8.**
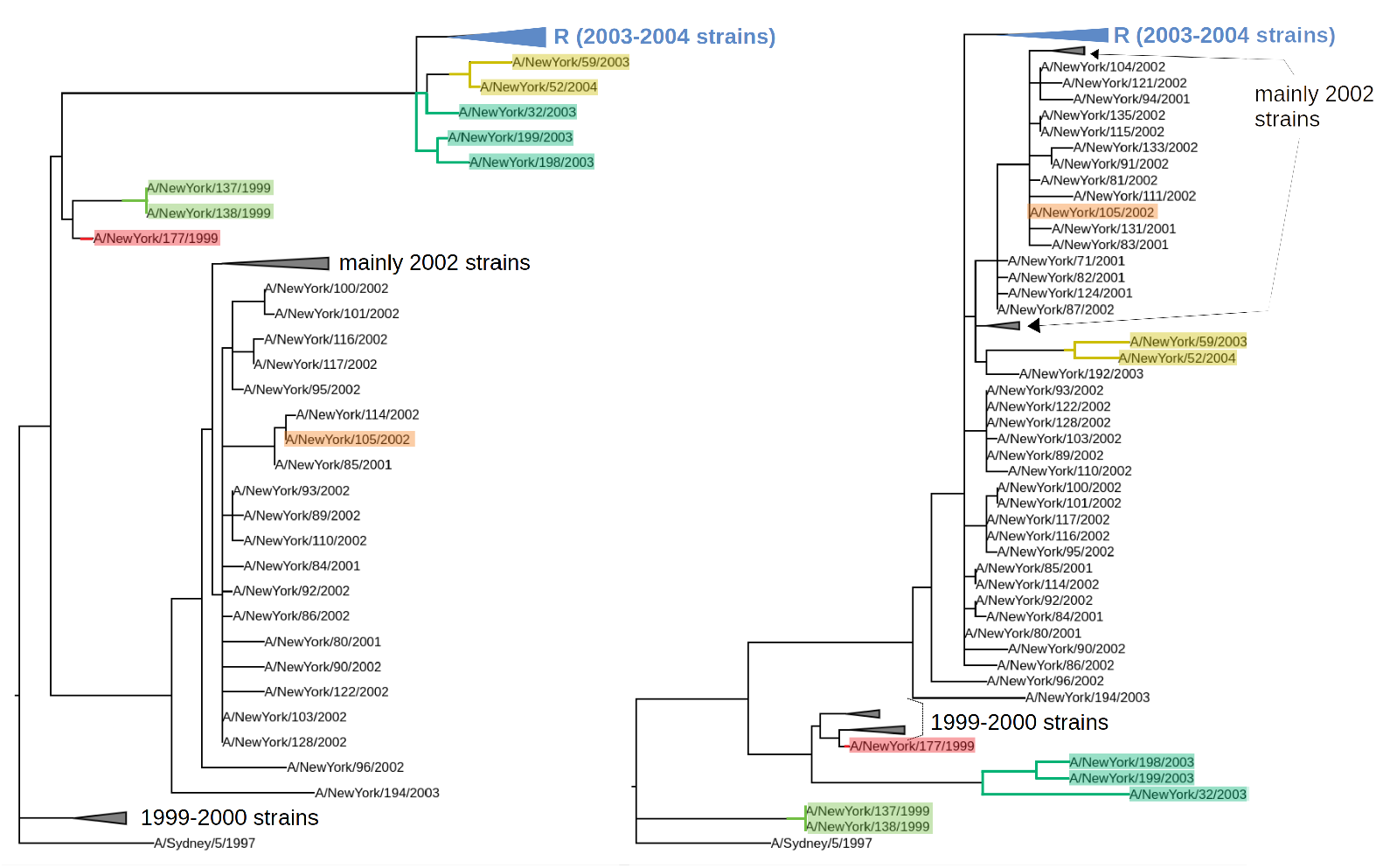
Segment trees for the 156 strains studied in [9]. **Left**: HA segment and **right**: NA segment. Some clades are collapsed for better visibility. The 6 MCCs corresponding to a reassortment found by our method (*γ* = 2) are highlighted. One of them involves a clade of 58 strains, shown collapsed and labelled as “`R`”. The remaining MCC contains all remaining strains as well as the root of both trees, and does not correspond to a reassortment. Previous studies ([9, 13]) only found reassortments for clades {A/New York/52/2004, A/New York/59/2003}, {A/New York/32/2003, A/New York/198/2003, A/New York/199/2003} and {A/New York/105/2003}. We also find clades {A/New York/137/2004, A/New York/138/2003}, {A/New York/177/1999} and {`R`}.

**Figure S 9.**
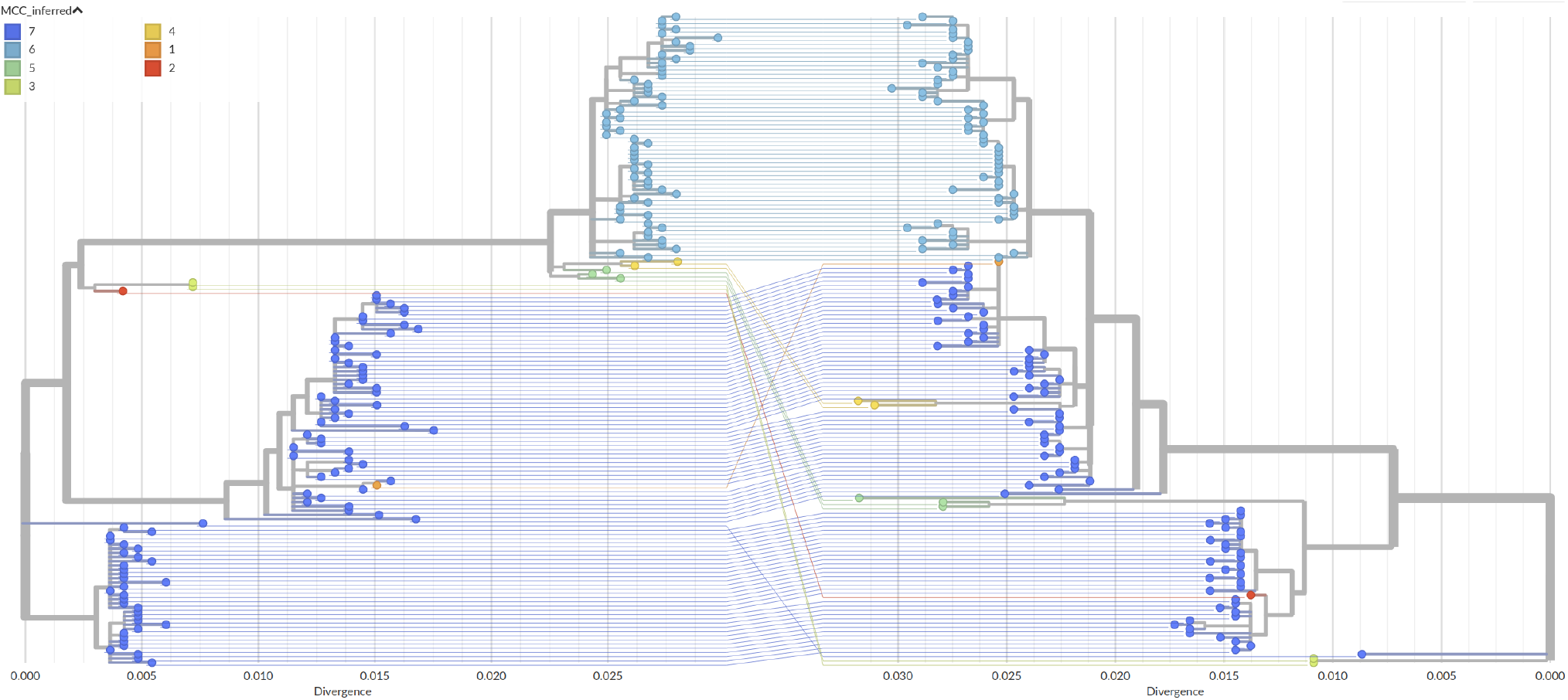
Segment trees and tanglegram for the 156 strains studied in [9]. Strains and branches are colored based on MCCs found by our method (*γ* = 2). MCC 7 does not correspond to a reassortment, as it contains the roots. The list of remaining MCCs is as follows: 1→{A/New York/105/2003}; 2→{A/New York/177/1999}; 3→{A/New York/137/2004, A/New York/138/2003}; 4→{A/New York/52/2004, A/New York/59/2003}; 5→{A/New York/32/2003, A/New York/198/2003, A/New York/199/2003}; 6→{`R`} (see Figure S8). Previous studies only found reassortments 1, 4 and 5 [9, 13]. Note that since MCCs have the same topology in the two trees, it is possible to completely disentangle lines of the same color in this plot.

**Figure S 10.**
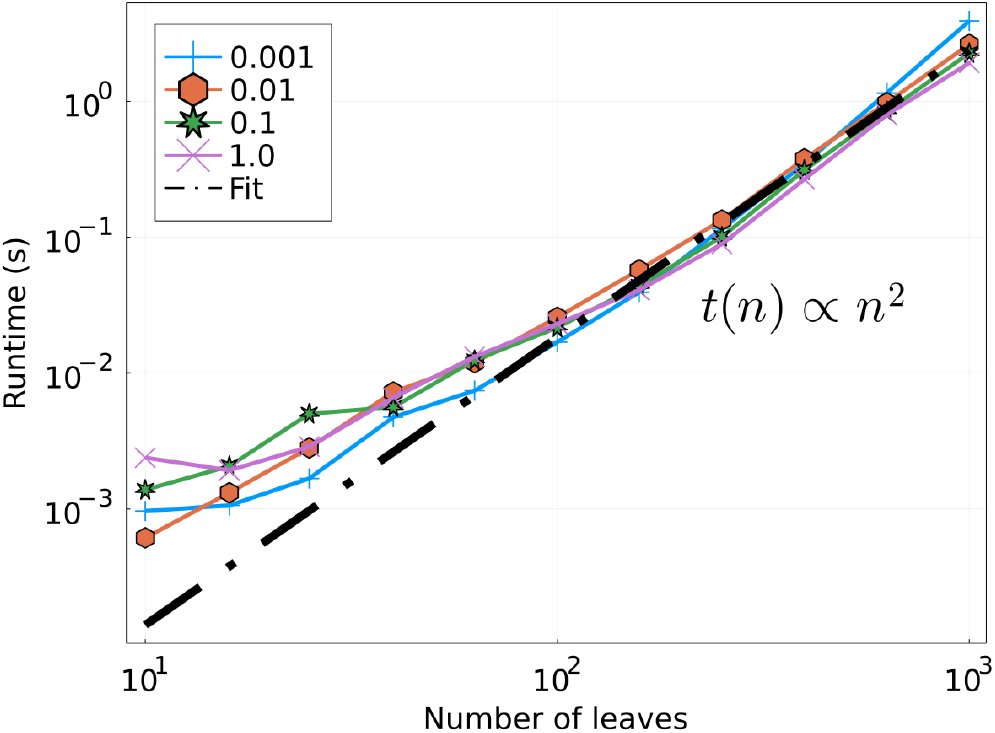
Runtime in seconds as a function of the number of leaves *L*, for different values of *ρ*, in log-log scale. Performed on a single CPU. The linear fit is done for *L >* 10^2^. As expected, runtime is quadratic in the number of leaves of the trees.

**Figure S 11.**
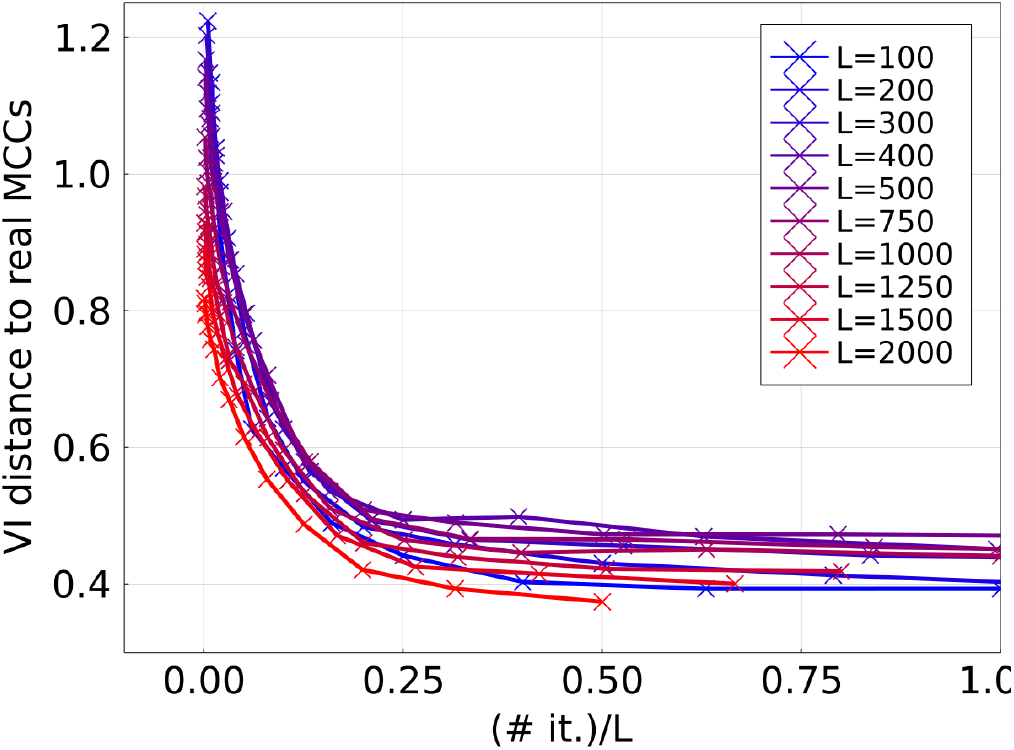
Estimation of the convergence of the algorithm for simulated ARGs of *L* leaves: VI distance to real MCCs as a function of the number of iterations of the SA optimization, scaled by *L*. The rhythm of convergence is the same for all curves, indicating that the number of iterations needed to reach convergence should be proportional to *L*.

**Figure S 12.**
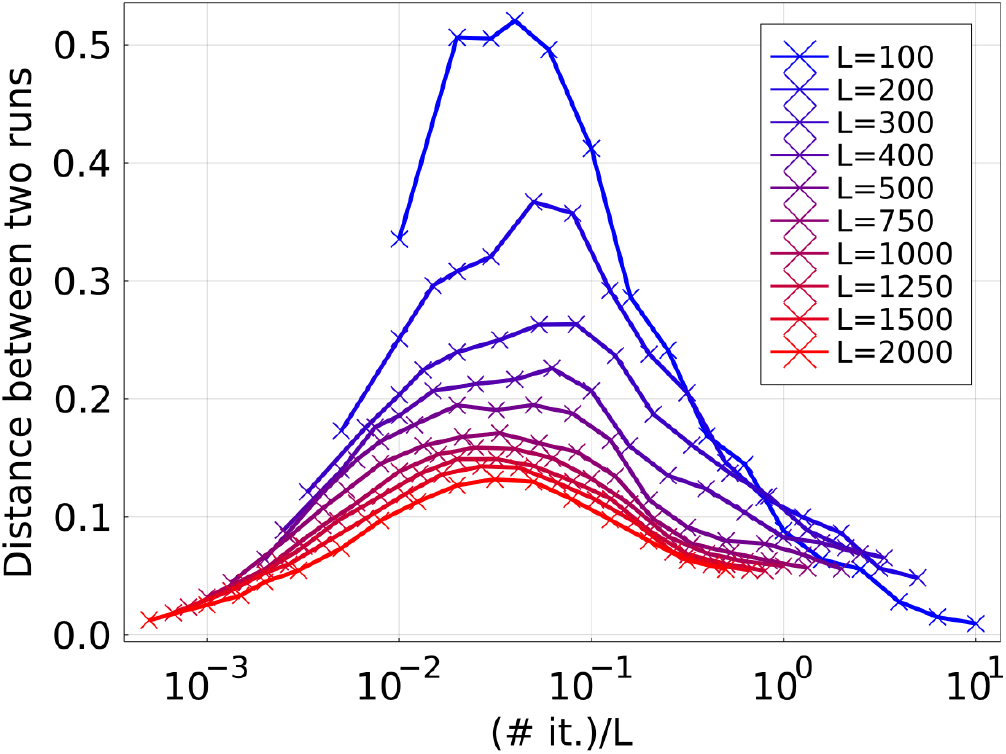
Estimation of the reproducibility of results for simulated ARGs of *L* leaves: VI distance between two independent runs as function of the number of iterations of the SA optimization, scaled by *L*. For a very low number of iterations, results are close to the naive MCCs, which are the starting point of the optimization. The distance between two runs is maximal for an intermediate number of iterations, and vanishes again as the optimization converges.

**Figure S 13.**
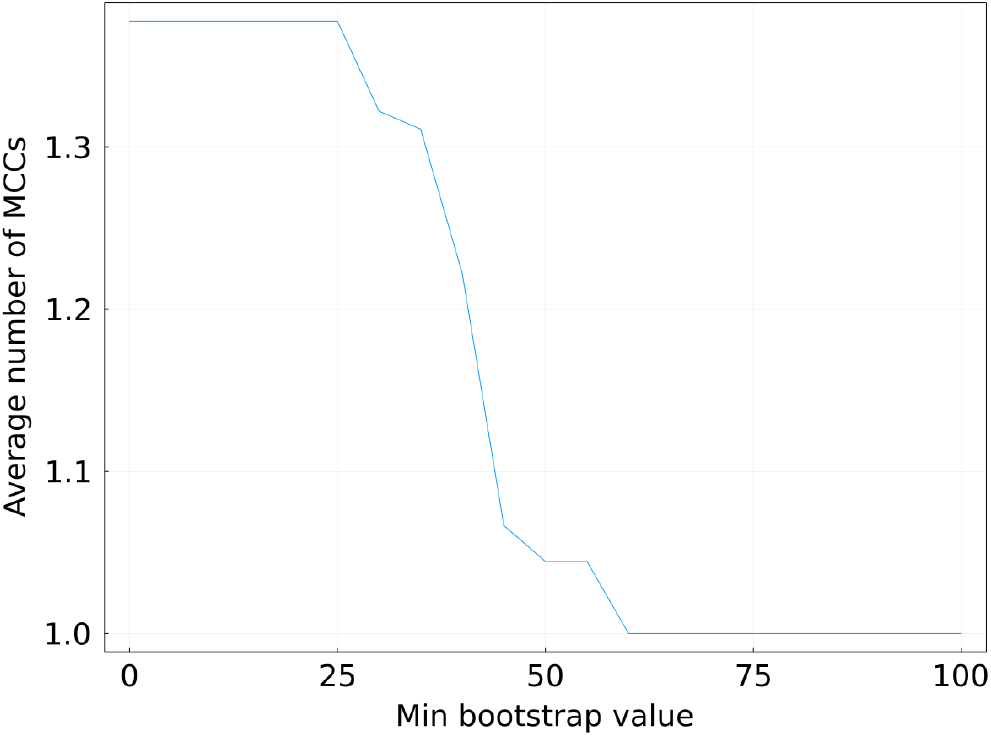
Evaluation of the robustness of our method with respect to the tree inference. On the *y*-axis: average number of MCCs obtained when applying the algorithm to two trees inferred from the same sequences. The average is performed using a set of 10 A/H3N2 HA alignments. Finding more than one MCC implies that some topological differences are introduced by the tree building process. On the *x*-axis: minimum bootstrap value required for branches. Branches with a smaller bootstrap value are removed from the trees. A minimum bootstrap of 75 is enough to guarantee an MCC inference robust to errors in the inference of the trees.

## Notes

### Competing Interest Statement

The authors have declared no competing interest.

## References

[1] Gavin J. D. Smith, Justin Bahl, Dhanasekaran Vijaykrishna, Jinxia Zhang, Leo L. M. Poon, Honglin Chen, Robert G. Webster, J. S. Malik Peiris, and Yi Guan. Dating the emergence of pandemic influenza viruses. Proceedings of the National Academy of Sciences, 106(28):11709–11712, July 2009. ISSN 0027-8424, 1091-6490. doi:10.1073/pnas.0904991106. URL http://www.pnas.org/lookup/doi/10.1073/pnas.0904991106.

[2] Yi Guan, Dhanasekaran Vijaykrishna, Justin Bahl, Huachen Zhu, Jia Wang, and Gavin J. D. Smith. The emergence of pandemic influenza viruses. Protein & Cell, 1(1):9–13, January 2010. ISSN 1674-800X. doi:10.1007/s13238-010-0008-z. URL https://www.ncbi.nlm.nih.gov/pmc/articles/PMC4875113/.

[3] Morgan N. Price, Paramvir S. Dehal, and Adam P. Arkin. FastTree: Computing Large Minimum Evolution Trees with Profiles instead of a Distance Matrix. Molecular Biology and Evolution, 26(7):1641–1650, July 2009. ISSN 0737-4038. doi:10.1093/molbev/msp077. URL https://doi.org/10.1093/molbev/msp077.

[4] Bui Quang Minh, Heiko A Schmidt, Olga Chernomor, Dominik Schrempf, Michael D Woodhams, Arndt von Haeseler, and Robert Lanfear. IQ-TREE 2: New Models and Efficient Methods for Phylogenetic Inference in the Genomic Era. Molecular Biology and Evolution, 02 2020. ISSN 0737-4038. doi:10.1093/molbev/msaa015. URL https://doi.org/10.1093/molbev/msaa015.msaa015.

[5] Alexandros Stamatakis. RAxML version 8: a tool for phylogenetic analysis and post-analysis of large phylogenies. Bioinformatics (Oxford, England), 30(9):1312–1313, May 2014. ISSN 1367-4811. doi:10.1093/bioinformatics/btu033.

[6] Alexey D. Neverov, Ksenia V. Lezhnina, Alexey S. Kondrashov, and Georgii A. Bazykin. Intrasubtype Reassortments Cause Adaptive Amino Acid Replacements in H3N2 Influenza Genes. PLOS Genetics, 10(1):e1004037, January 2014. ISSN 1553-7404. doi: 10.1371/journal.pgen.1004037. URL https://journals.plos.org/plosgenetics/article?id=10.1371/journal.pgen.1004037. Publisher: Public Library of Science.

[7] Mara Villa and Michael Lässig. Fitness cost of reassortment in human influenza. PLOS Pathogens, 13(11):e1006685, November 2017. ISSN 1553-7374. doi: 10.1371/journal.ppat.1006685. URL https://journals.plos.org/plospathogens/article?id=10.1371/journal.ppat.1006685. Publisher: Public Library of Science.

[8] Nicola F. Müller, Ugnė Stolz, Gytis Dudas, Tanja Stadler, and Timothy G. Vaughan. Bayesian inference of reassortment networks reveals fitness benefits of reassortment in human influenza viruses. Proceedings of the National Academy of Sciences, 117(29):17104–17111, July 2020. ISSN 0027-8424, 1091-6490. doi:10.1073/pnas.1918304117. URL https://www.pnas.org/content/117/29/17104. Publisher: National Academy of Sciences Section: Biological Sciences.

[9] Edward C. Holmes, Elodie Ghedin, Naomi Miller, Jill Taylor, Yiming Bao, Kirsten St George, Bryan T. Grenfell, Steven L. Salzberg, Claire M. Fraser, David J. Lipman, and Jeffery K. Taubenberger.Whole-Genome Analysis of Human Influenza A Virus Reveals Multiple Persistent Lineages and Reassortment among Recent H3N2 Viruses. PLOS Biology, 3(9):e300, July 2005. ISSN 1545-7885. doi:10.1371/journal.pbio.0030300. URL https://journals.plos.org/plosbiology/article?id=10.1371/journal.pbio.0030300. Publisher: Public Library of Science.

[10] Martha I. Nelson, Cécile Viboud, Lone Simonsen, Ryan T. Bennett, Sara B. Griesemer, Kirsten St George, Jill Taylor, David J. Spiro, Naomi A. Sengamalay, Elodie Ghedin, Jeffery K. Taubenberger, and Edward C. Holmes. Multiple Reassortment Events in the Evolutionary History of H1N1 Influenza A Virus Since 1918. PLOS Pathogens, 4(2):e1000012, February 2008. ISSN 1553-7374. doi:10.1371/journal.ppat.1000012. URL https://journals.plos.org/plospathogens/article?id=10.1371/journal.ppat.1000012. Publisher: Public Library of Science.

[11] Raul Rabadan, Arnold J. Levine, and Michael Krasnitz. Non-random reassortment in human influenza A viruses. Influenza and Other Respiratory Viruses, 2 (1):9–22, 2008. ISSN 1750-2659. doi:10.1111/j.1750-2659.2007.00030.x. URL https://onlinelibrary.wiley.com/doi/abs/10.1111/j.1750-2659.2007.00030.x.eprint: https://onlinelibrary.wiley.com/doi/pdf/10.1111/j.1750-2659.2007.00030.x.

[12] U. Chandimal de Silva, Hokuto Tanaka, Shota Nakamura, Naohisa Goto, and Teruo Yasunaga. A comprehensive analysis of reassortment in influenza A virus. Biology Open, 1(4):385–390, February 2012. ISSN 2046-6390. doi:10.1242/bio.2012281. URL https://www.ncbi.nlm.nih.gov/pmc/articles/PMC3509451/.

[13] Niranjan Nagarajan and Carl Kingsford. GiRaF: robust, computational identification of influenza reassortments via graph mining. Nucleic Acids Research, 39(6):e34–e34, March 2011. ISSN 0305-1048. doi:10.1093/nar/gkq1232. URL https://doi.org/10.1093/nar/gkq1232.

[14] Alisa Yurovsky and Bernard M E Moret. FluReF, an automated flu virus reassortment finder based on phylogenetic trees. BMC Genomics, 12(2):S3, July 2011. ISSN 1471-2164. doi:10.1186/1471-2164-12-S2-S3. URL https://doi.org/10.1186/1471-2164-12-S2-S3.

[15] Victoria Svinti, James A. Cotton, and James O. McInerney. New approaches for unravelling reassortment pathways. BMC Evolutionary Biology, 13(1):1, January 2013. ISSN 1471-2148. doi:10.1186/1471-2148-13-1. URL https://doi.org/10.1186/1471-2148-13-1.

[16] Ugnė Stolz, Tanja Stadler, Nicola F. Müller, and Timothy G. Vaughan. Joint inference of migration and reassortment patterns for viruses with segment genomes. Technical report, May 2021. URL https://www.biorxiv.org/content/10.1101/2021.05.15.442587v1. xCompany: Cold Spring Harbor Laboratory Distributor: Cold Spring Harbor Laboratory Label: Cold Spring Harbor Laboratory Section: New Results Type: article.

[17] S. Kirkpatrick, C. D. Gelatt, and M. P. Vecchi. Optimization by Simulated Annealing. Science, 220(4598):671–680, May 1983. doi:10.1126/science.220.4598.671. URL https://www.science.org/lookup/doi/10.1126/science.220.4598.671. Publisher: American Association for the Advancement of Science.

[18] Gabriel Cardona, Francesc Rosselló, and Gabriel Valiente. Extended Newick: it is time for a standard representation of phylogenetic networks. BMC Bioinformatics, 9(1):532, December 2008. ISSN 1471-2105. doi:10.1186/1471-2105-9-532. URL https://doi.org/10.1186/1471-2105-9-532.

[19] Marina Meilă. Comparing clusterings—an information based distance. Journal of Multivariate Analysis, 98(5):873–895, May 2007. ISSN 0047-259X. doi:10.1016/j.jmva.2006.11.013. URL https://www.sciencedirect.com/science/article/pii/S0047259X06002016.

[20] Niranjan Nagarajan and Carl Kingsford. Uncovering Genomic Reassortments among Influenza Strains by Enumerating Maximal Bicliques. In 2008 IEEE International Conference on Bioinformatics and Biomedicine, pages 223–230, November 2008. doi:10.1109/BIBM.2008.78.

[21] John P. Huelsenbeck and Fredrik Ronquist. MRBAYES: Bayesian inference of phylogenetic trees. Bioinformatics, 17(8):754–755, August 2001. ISSN 1367-4803. doi: 10.1093/bioinformatics/17.8.754. URL https://doi.org/10.1093/bioinformatics/17.8.754.

[22] Lam-Tung Nguyen, Heiko A. Schmidt, Arndt von Haeseler, and Bui Quang Minh. IQTREE: A Fast and Effective Stochastic Algorithm for Estimating Maximum-Likelihood Phylogenies. Molecular Biology and Evolution, 32(1):268–274, 11 2014. ISSN 0737-4038. doi:10.1093/molbev/msu300. URL https://doi.org/10.1093/molbev/msu300.

[23] R. R. Hudson and N. L. Kaplan. Deleterious Background Selection with Recombination. Genetics, 141(4):1605–1617, December 1995. ISSN 0016-6731. URL https://www.ncbi.nlm.nih.gov/pmc/articles/PMC1206891/.

[24] W. G. Hill and Alan Robertson. The effect of linkage on limits to artificial selection. Genetics Research, 8(3):269–294, December 1966. ISSN 1469-5073, 0016-6723. doi:10.1017/S0016672300010156. URL https://www.cambridge.org/core/journals/genetics-research/article/effect-of-linkage-on-limits-to-artificial-selection/5CCFE11C1F8108242ED02AEC8BA5DD50.

[25] R. A. Neher and B. I. Shraiman. Genetic draft and quasi-neutrality in large facultatively sexual populations. Genetics, 188(4):975–996, August 2011. ISSN 1943-2631 0016-6731. doi:10.1534/genetics.111.128876.

[26] Gytis Dudas, Trevor Bedford, Samantha Lycett, and Andrew Rambaut. Reassortment between Influenza B Lineages and the Emergence of a Coadapted PB1–PB2–HA Gene Complex. Molecular Biology and Evolution, 32(1):162–172, January 2015. ISSN 0737-4038. doi: 10.1093/molbev/msu287. URL https://doi.org/10.1093/molbev/msu287.

[27] James Hadfield, Colin Megill, Sidney M Bell, John Huddleston, Barney Potter, Charlton Callender, Pavel Sagulenko, Trevor Bedford, and Richard A Neher. Nextstrain: real-time tracking of pathogen evolution. Bioinformatics, 34(23):4121–4123, 05 2018. ISSN 1367-4803. doi: 10.1093/bioinformatics/bty407. URL https://doi.org/10.1093/bioinformatics/bty407.

